# HSF1-dependent, long-range directional *HSPA1* gene motion to nuclear speckles is coupled to DDX39B condensate dynamics

**DOI:** 10.64898/2026.01.07.698255

**Authors:** Gabriela Andrea Hernandez Gonzalez, Purnam Ghosh, Jiah Kim, Saireet Misra, Pradeep Kumar, Pankaj Chaturvedi, Neha Chivukula Venkata, Andrew S. Belmont

## Abstract

Approximately 5-10% of the genome, enriched ∼10-fold in the most highly active genes, positions near deterministically within several hundred nm of nuclear speckles (NS). This includes the *HSPA1* locus, among the most strongly NS-associated genes. Here we address how *HSPA1* endogenous genes and transgenes stably position adjacent to the nuclear speckle periphery, facilitating their gene expression amplification. Live-cell imaging demonstrates a transcription-dependent speckle-anchoring of *HSPA1* genes. Removal of this anchoring reveals sustained, long-range, curvilinear oscillations of the HSPA1 genes away from and back to nuclear speckles or between nuclear speckles. Curvilinear transgene movements are dependent on the transcription factor HSF1, largely independent of nuclear actin or cohesin, but unexpectedly spatially and temporally correlated with DDX39B condensate dynamics and dependent on DDX39B protein levels. We propose that stable nuclear speckle association of *HSPA1* results from combining metastable speckle anchoring with constitutive, restorative long-range chromosome movements driven by DDX39B condensate dynamics.

## Introduction

The mammalian genome folds nonrandomly within the nucleus with differential positioning of chromosome regions relative to several nuclear locales highly correlated with their differential gene expression and DNA replication timing^1–8^. One such nuclear locale are nuclear speckles (NS), the light microscopy equivalent of the Interchromatin Granule Clusters (IGCs) visualized by electron microscopy^9–11^. The preferential localization of a subset of active genes adjacent to NS was described decades ago^9^. Recent genome-wide sequencing-based measurements using TSA-seq revealed that the top 5% of the genome closest to NS show a mean NS distance less than ∼300 nm, with >90-95% of alleles of representative examples positioned within ∼500 nm^12^. These speckle-associated genomic regions represent local gene expression “hot-zones”, highly enriched for the most highly expressed genes, super-enhancers, and epigenetic markers associated with active chromatin^12,13^. A physiological link between NS association and gene expression was suggested by the live-cell demonstration of increased gene expression of *HSPA1* heat shock genes after NS contact^14,15^; a similar phenomenon of gene expression amplification after NS contact was also inferred from fixed cell data for several genes flanking the *HSPA1* locus^15^, the *HSPH1* gene^16^, and several hundred p53-responsive genes^17^. Additionally, most genes significantly downregulated after elimination of NS by SON and SRRM2 double knockdown are those positioned very close to NS, suggesting that these genes also may show gene expression amplification with NS contact^18,19^.

A fundamental question then becomes how many gene loci are stably positioned adjacent to NS. Outside of early G1, during which long-range chromosome movements can be observed, dynamics of individual mammalian chromosome loci generally fit a “constrained diffusion” model with apparently diffusive short-range chromosome movements constrained over longer times within a radius of ∼0.5 microns^20–22^. Indeed, recent measurements of chromatin anomalous diffusion of chromatin has led to the prediction that whereas genomic elements within several hundred nm of each other would rapidly come into contact with each other, estimated times for contact of genomic elements separated by micron-scale distances would be “impossibly long” ^23^. Thus, common models postulate that chromosomes diffuse randomly within early G1 mammalian nuclei, bind to or nucleate specific compartments/bodies, and then remain relatively immobile throughout the remaining interphase, with substantial movement of chromosomes between nuclear compartments not occurring until progression through the next mitosis^22^.

We previously used *HSPA1* genes and transgenes as a model system to probe gene positioning relative to NS^14,15,24,25^. We demonstrated a strong association of BAC *HSPA1* transgenes with NS^14,15,24^, reproducing the unusually strong NS association observed for the endogenous *HSPA1* chromosome locus both before and after heat shock^15,16^. Using promoter swap experiments, we observed that the increased NS association after heat shock of randomly integrated *HSPA1*A plasmid arrays required the 5’ promoter and proximal regulatory sequences of *HSPA1A* but not the gene body sequence itself^25^; we also demonstrated decreased speckle association after transcriptional inhibition^25,26^. At that time, we suggested that certain cis regulatory promoter sequences might direct specific chromatin and/or RNP transcript modifications leading to NS association.

Here we reexamine how the *HSPA1* locus stably associates with NS. Exploiting live-cell imaging, we demonstrate how stable NS association is separable into the mechanistically distinct processes of stable anchoring versus micron-scale, curvilinear movements which move alleles away from but then back towards NS and between neighboring NS. By removing this stable anchoring, our results reveal a non-diffusive, constitutive, transcription factor and nuclear condensate dependent mechanism for moving genes long-distance and directionally within nuclei over a several minute timescale.

## Results

### Experimental System

Previously, we discovered the targeting of *HSPA1* BAC transgenes to NS at 37°C, which increased with heat shock. More specifically, an ∼2 Mbp, 10-copy BAC transgene array was attached at very high frequency at one or more BAC copies to NS with heat shock inducing both the decondensation of the BAC array and its wrapping around the NS circumference^24^. Additionally, we observed *HSPA1* nascent mRNAs accumulating in the adjacent NS^24,25^, thus classifying *HSPA1* as a “Type 1” NS associated gene locus different from the more frequent “Type 2” NS associated genes whose nascent transcripts accumulate nearby but outside the NS^9^. A survey of ∼20 highly active genes revealed approximately half to be NS associated but only a couple to be Type 1 associated genes^9^.

To determine genome-wide how many endogenous genes associate specifically with NS, we developed TSA-seq, which provides a quantitative estimate of the mean distance of genes to NS^12^. Interestingly, several heat shock gene loci including *HSPA1* mapped in the top ∼1% of SON TSA-seq scores^16^. We confirmed this close NS association of nearly all *HSPA1* alleles across multiple cell lines using FISH^16^.

These results place the *HSPA1* locus’s nuclear speckle association among the strongest genome-wide making it an excellent choice for dissection of the mechanisms underlying gene association with NS. Here we used two cell lines previously engineered for live-cell imaging (**Fig. 1**).

**Figure 1.**
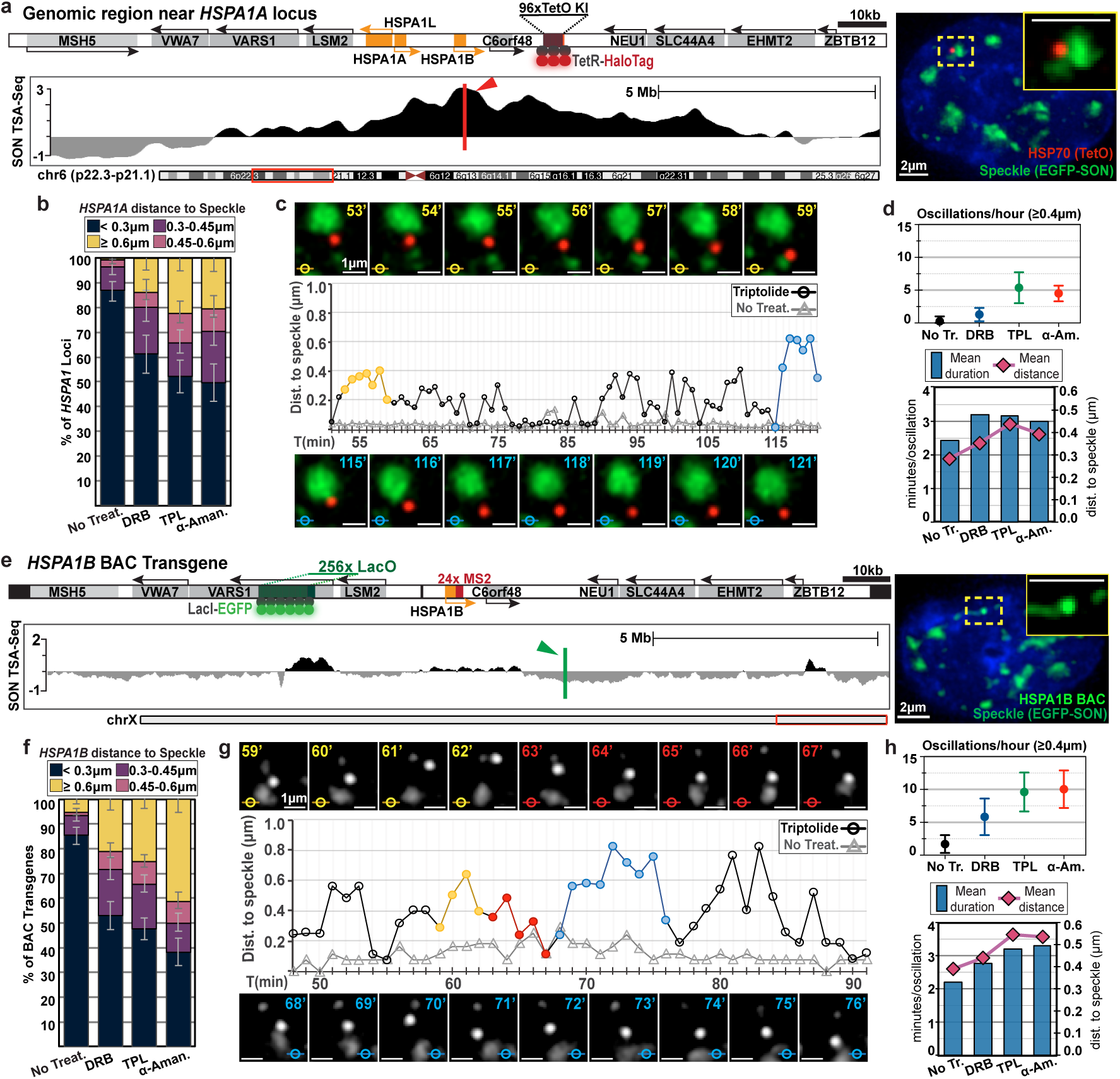
Transcriptional inhibition increases gene distance to nuclear speckles and induces dynamic gene oscillations. (**a & e**), Top left panel-Diagrams of the human *HSPA1* locus with HSPA1 (yellow) and flanking (grey) genes in the HCT116 “96-8 B2” cell line (**a**) and the integrated *HSPA1B* BAC transgene in the CHOK1 “C7MCP” cell line (**e**). Positions of the 96-mer TetO repeats bound by TetR-HaloTag (red) and 256-mer LacO repeats bound by LacI-EGFP (green) are indicated. Right panel-Representative images show HSPA1 locus **(a)** (red) or *HSPA1B* BAC **(e)** (bright green and nuclear speckles (NS) (green, EGFP-SON), with DNA (DAPI, blue). Bottom panel-SON TSA-seq tracks show genomic microscopic proximity to NS (**a**: human chr6; **e**: CHO chrX); high enrichment scores (black peaks) indicate chromosome regions with mean distances to NS of just several hundred nm. The *HSPA1* locus (**a**) BAC integration sites (**e**) are marked (red/green arrows). (**b & f)**, Histograms show fraction of *HSPA1* loci (**b**) or *HSPA1B* BAC transgenes (**f**) at defined distances from NS in fixed cells before and after treatment with transcriptional inhibitors at 37°C. Data pooled from three replicates: endogenous *HSPA1* (n=145–270) and BAC transgene (n=283–480) cells. Error bars represent 95% confidence intervals calculated using the Wilson Score method (see Methods for details). Overlapping error bars reflect sampling uncertainty in proportion estimates. (**c & g**), Live-cell imaging (1 frame/min) reveals continuous, bidirectional movement of gene loci relative to NS after inhibitor treatment. Consecutive frames show oscillations between NS (top, yellow/red timestamp) and to/from a single speckle (bottom, blue). Middle panels show plots of gene–NS distances versus time, with color-coded points indicating corresponding color-coded frames shown above and below as images; grey traces show similar gene-NS distance plots from untreated control cells. (**d & h)**, Top panel-Mean number of oscillations per hour (events with gene-speckle separation of ≥ 0.4 µm) (mean ± SD; Total nuclei counted (d) n=7, 5, 6, 5 and (h) n= 9, 8, 10, 10 per treatment). Bottom panel-bar graphs show mean duration of individual oscillations (minutes) and mean maximum locus-NS distance (µm) per oscillation following inhibitor treatment. Total oscillation counts: (**d**), n= 54, 55, 123, 123 and (**h**), n= 45, 152, 234 and 265 for untreated (No Treat.), DRB, triptolide (TPL) and alpha-amanitin (α-Aman.) treated cells, respectively.

To track the endogenous locus, we used a human colon carcinoma HCT116 cell line in which a 96-mer Tet operator (TetO) array was inserted ∼16 kb downstream of one of the endogenous *HSPA1B* gene alleles (**Fig. 1a**, left panel, top)^27^. We then used CRISPR to tag the endogenously expressed nuclear speckle marker, SON, with EGFP to create cell line “HCT116 96-8 B2”, followed by plasmid transfection to create a mixed population of cell clones stably expressing a Tet Repressor (TetR)-HaloTag fusion protein (**Fig. 1a**, right panel). The *HSPA1* locus, containing homologous *HSPA1A*, *HSPA1B*, and *HSPA1L* genes, is positioned within a large SON TSA-seq peak flanked by several SON TSA-seq subpeaks within a broad, several Mbp ridge of elevated SON TSA-seq (**Fig. 1a**, left panel, bottom).

To track a *HSPA1* BAC transgene, we used the previously described C7MCP CHO cell clone stably expressing GFP-Lac repressor (GFP-LacR) and containing an estimated 1-3 copies of an *HSPA1B* BAC transgene^15^. This *HSPA1B* BAC containing ∼200 kb of the human *HSPA1* locus has two of the three *HSPA1* genes (*HSPA1A* and *HSPA1L*) deleted and is tagged with a 256-mer Lac operator (LacO) array inserted upstream of the *HSPA1B* gene and a 24-mer MS2 repeat inserted into the *HSPA1B* 3’ UTR (**Fig. 1e**, left panel, top)^14^. To label NS, this clone was stably transformed with an EGFP-SON BAC transgene (**Fig. 1e**, right panel) and, after this, with an mCherry-MS2 binding protein (MS2BP) plasmid transgene^15^. Long DNA sequencing revealed the insertion of the BAC transgene within a SON TSA-seq valley located several Mbp away both 5’ and 3’ from the nearest large SON TSA-seq peak (**Fig. 1e**, left panel, bottom).

Fixed cell measurements show nearly as strong NS association of the BAC transgene as the endogenous *HSPA1* locus in human cells under normal growth conditions (**Fig. 1b&f, SFig. 1a-b**, “No treat”).

### Transcriptional inhibition destabilizes *HSPA1* NS anchoring resulting in sustained HSPA1 gene movements to and from and between NS

Previously, we showed a near complete or partial elimination of NS association after heat shock of a HSPA1 plasmid transgene array using alpha-amanitin^25^ or DRB (5,6-dichloro-1-beta-D-ribofuranosylbenzimidazole), respectively to inhibit transcription^28^. Here we show a significantly reduced NS association of both the endogenous *HSPA1* locus and the integrated *HSPA1B* BAC, as measured in fixed cells, after inhibiting RNA polymerase II (RNAP II) by three different inhibitors acting through different mechanisms (**Fig. 1b&f**, **SFig 1a-b**): DRB, inhibiting RNA polymerase II (RNAP II) transcriptional pause release, triptolide (TPL), inhibiting RNAP II transcriptional initiation, and alpha-amanitin, which both blocks RNAP II elongation and degrades RNAP II under the conditions used ^29^. All three inhibitors significantly reduced *HSPA1* NS association both before and after heat shock (HS) at 42°C for 30 min; alpha-amanitin produced the largest reduction and DRB the smallest (**Fig. 1b&f**, **SFig. 1a-b**). Effects were similar for both the endogenous *HSPA1* locus and *HSPA1B* BAC transgene.

Whereas fixed cell data suggested transcriptional inhibition produced loss of HSPA1 speckle association in only a fraction of cells, live cell movies unexpectedly revealed that transcriptional inhibition caused dynamic detachment and then reattachment of HSPA1 genes and transgenes from NS in essentially all cells over time (Fig. 1c&g, Movie 1). Distance histograms derived from live-cell imaging essentially reproduced the fixed cell data both before and after TPL inhibition (**SFig. 1c**), demonstrating minimal phototoxicity.

To better visualize and measure gene and transgene dynamics relative to adjacent NS, we first projected the optical section stacks from each time along the optical (z) axis. We then used a 2D cross-correlation alignment to correct for the translation and/or rotation of nuclei between adjacent time-points. After this we measured the 2D distance of the *HSPA1* locus to the nearest NS edge for each time frame (**Fig. 1**, **SFig. 1**).

In untreated cells, both the endogenous *HSPA1* locus and the BAC transgene remain stably associated with NS most of the time, with rare detachments followed by quick reattachments (**Fig. 1c&g**, **SFig. 1f; Movie 1**, left panels). We define an “oscillation” as any movement of a speckle-associated locus away from a NS followed by its return and reattachment to the same or a different NS (See Methods). Inhibition with any of the three RNAP II inhibitors significantly increased the frequencies of these oscillatory movements for both the endogenous locus and the BAC transgene, with the greatest effects observed for triptolide and alpha-amanitin (**Fig. 1d&h, SFig. 1d**; **Movies 1&2**).

This increase in oscillation frequencies was especially pronounced for the BAC transgene, which showed twice as many long-range oscillations (≥0.4 microns from the NS) compared to the endogenous HSPA1 locus **(Fig. 1d&h)**. Consistent with results from fixed cells, the BAC transgene showed a slightly higher average distance from NS per oscillation (**Fig. 1d & 1h**, **SFig. 1e**). While the frequencies of movements from one NS to another were similar for both the endogenous and transgene loci (**SFig. 1e**), we noticed an increase in movements between more distant NS for the BAC transgene (data not shown) We speculate that the more limited range of motion from NS seen for the endogenous locus is due to additional anchoring of adjacent chromosome regions to the same NS via neighboring NS-associated domains flanking the endogenous gene but not the transgene **(Fig. 1a&e, left panels, bottom).**

In summary, we conclude that transcriptional inhibition leads to a loss of *HSPA1* gene and transgene NS anchoring, resulting in a large increase in oscillatory dynamics on and off nuclear speckles and between NS.

### BAC transgene movements retrace similar curvilinear paths to and from and between nuclear speckles

Our further analysis focused on the BAC transgene system due to its ∼50% higher frequency of oscillations, larger maximum distance excursions from NS, and longer-distance movements between NS. Live cell movies revealed that genes and transgenes followed approximately curvilinear trajectories during oscillations and movements between nuclear speckles after transcriptional inhibition **(Movie 2).**

To better visualize the paths of BAC transgene movements, we next applied a temporal color-coded projection on our 2D time-lapse sequences (**Fig. 2**). This technique assigns distinct colors from lookup table (LUT) to sequential time frames, with early time points displayed in colors from one end of the LUT and later frames progressively transitioning toward the opposite end. (**Fig. 2a & b**). To display unidirectional movements- or lack thereof-we utilized the Spectrum LUT (**Fig. 2a**, **Movie 2**). This visualization reveals transgene movement and direction over time while highlighting stationary regions whose colors combine to produce white coloration. We used the Blue-Magenta-Red (BMR) and Cyan-Green-Yellow (CGY) LUTs (**Fig. 2b**, **Movie 2**) to display transgene movement away from and back to nuclear speckles, respectively. Beyond indicating transgene direction during oscillations, these three-color LUTs are complementary in the color spectrum, enabling us to overlay/merge these temporally-coded images while distinguishing their distinct paths. Where these trajectories overlap, the complementary colors combine to produce different colors up to white coloration.

**Figure 2.**
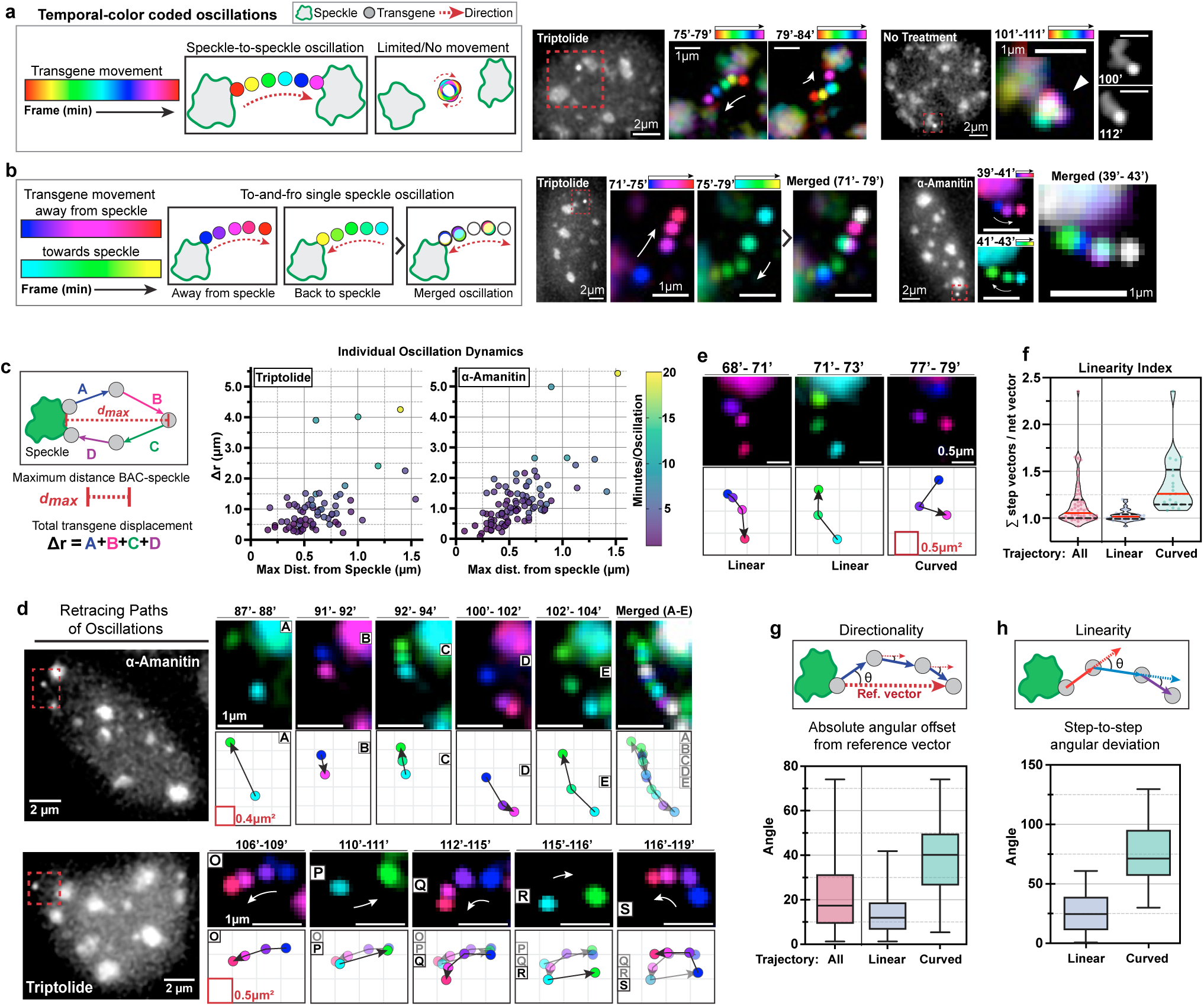
*HSPA1B* BAC transgene oscillations follow curvilinear, directional trajectories to and from or between speckles. **(a)** Left: Schematic of temporal color-coded projection using Spectrum LUT (left). Right: Representative examples of BAC transgene movement after triptolide (TPL) treatment versus control showing white coloration from overlapping frames (right). (**b)** Left: Temporal projections using complementary BMR and CGY LUTs showing movement away from and back to nuclear speckles (NS), respectively. Right: Examples after TPL or α-amanitin treatment with overlapping paths producing intermediate colors. (**c)** Left: Diagram showing measurements per oscillation: Maximum distance from NS (d_max_; µm) is the largest separation between transgene and nearest NS edge; total displacement (Δr) is the sum of frame-to-frame step distances. Right: Individual oscillation scatterplot statistics for *HSPA1B* BAC transgene after TPL or α-amanitin treatment. Color indicates duration (minutes; color bar) at 1 frame/min acquisition. n= 68 and 90 oscillations from 3 nuclei after TPL or α-amanitin treatment, respectively. (**d)** Two examples showing transgene bidirectional movements along approximately the same linear path through multiple to and fro movements using both temporal projection of the color-coded images and mapping of transgene positions to a spatial grid. Top: α-amanitin treatment example showing retracing of transgene path throughout multiple oscillations. Bottom: TPL treatment example showing transgene returning to similar positions and path retracing during continuous oscillations. (**e-h**) Statistics summarizing the extent of linearity and directionality of trangene movements. (**e**) Two examples of “linear” trajectories versus one example of nonlinear “curved” path; (**f**) Violin plots showing linearity index (ratio of sum of all step lengths to straight line distance between starting and end points) for all transgene trajectories (n=51), classified as “linear” (n=32), and “curved” (n=19). Linear trajectories displayed significantly lower median linearity index (1.017) compared to curved (1.261; Mann-Whitney U = 21, p < 0.0001). All trajectories: median = 1.053 (range: 1.000–2.357). (**g)** Angular measurements for BAC locus directionality. Top: cartoon showing blue step vectors (frame-to-frame displacements) and red dashed reference vector (start-to-end trajectory line); absolute angular offset measured between each step vector and reference vector. Bottom: box plots of absolute angular offsets for all step vectors (n = 111), linear (n = 69), and curved (n = 42) trajectories. Rayleigh test revealed highly significant directional clustering (R = 0.954, mean direction = 22.7°, p < 2.2 × 10⁻¹). Median [IQR]: All = 17.36° [9.315–31.51°], Linear = 11.90° [6.624–18.90°], Curved = 40.13° [26.44–49.80°]. Mean ± SD: All = 22.98° ± 17.75°, Linear = 13.61° ± 8.987°, Curved = 38.38° ± 17.89°. **(h)** Top: cartoon showing angular deviation between consecutive step vectors. Bottom: box plots of step-to-step angular deviations for linear (n = 37 step pairs) and curved (n = 23 step pairs) trajectories. Both groups showed significant deviations from 0° (linear: 26.18° ± 16.54°, t(36) = 9.63, p < 0.0001; curved: 73.20° ± 26.28°, t(22) = 13.36, p < 0.0001). Curved trajectories displayed 2.8-fold greater angular variability.

Whereas NS remain relatively stable in position, appearing in white, the transgenes exhibit curvilinear movement. Temporal projections over several minutes corresponding to unidirectional movements between NS (**Fig. 2a, right panel**) or away from and back to a single NS (**Fig. 2b, right panel; Fig. 2d**) reveal curvilinear BAC transgene trajectories, indicating an underlying non-diffusive mechanism. These curvilinear trajectories show significantly longer excursion distances than the smaller positional fluctuations observed over longer periods for speckle-associated genes under control conditions without transcriptional inhibition (S**Fig. 2 a&b**, “no treatment”; **Fig. 2c**, right panel).

In addition to maximum separation distance between gene locus and speckle (d_max), we extracted key quantitative metrics from locus position changes to characterize oscillations: step size (distance between positions in consecutive frames, μm)—used as a measure of overall gene mobility—and total transgene displacement (Δr) per oscillation (sum of steps within an oscillation) **(Fig. 2c**). These parameters, combined with oscillation frequency, distinguish BAC-driven oscillations from those resulting from speckle movement or nucleation.

A total of 51 gene locus trajectories were analyzed and classified into linear (n = 32) and curved (n = 19) groups based on average angular offset from the reference vector thresholds (≤25° and >25°, respectively) (**Fig. 2e**). Linear trajectories displayed a median linearity index of 1.017, while curved trajectories exhibited a significantly higher median linearity index of 1.261 (Mann-Whitney U = 21, p < 0.0001), validating the robustness of the angular offset-based classification scheme (**Fig. 2f**). Pooled analysis of all step vector absolute angular offsets (n = 111) revealed highly significant clustering around a preferred direction (Rayleigh test: R = 0.954, mean direction = 22.7°, p < 2.2 × 10^−1^□), strongly rejecting the null hypothesis of uniform angular distribution (**Fig. 2g**). This result provides compelling evidence that gene locus trajectories exhibit directional rather than random movement patterns. Both linear and curved trajectories exhibited significant step-to-step angular deviations (**Fig. 2h**). Linear trajectories showed mean deviations of 26.18° ± 16.54° (n = 37; t(36) = 9.63, p < 0.0001), while curved trajectories displayed substantially larger deviations of 73.20° ± 26.28° (n = 23; t(22) = 13.36, p < 0.0001), demonstrating approximately 2.8-fold greater angular variability in curved trajectories. Given that we are unable to correct for observed changes in nuclear shape between time points, these estimates are lower bounds on the actual trajectory linearity.

We previously described two similar examples of chromosome long-range, curvilinear movements: a heterochromatic plasmid array moving from the nuclear periphery towards the interior after tethering the VP16 acidic activator domain via a 256mer lac operator repeat^30^, and a multi-copy *HSPA1A* plasmid array moving toward nuclear speckles after heat shock^14^. In both systems, movement occurred asynchronously and typically once per observation period, with rare instances of retrograde movements.

In contrast, these *HSPA1B* BAC transgene oscillations to and from nuclear speckles after transcriptional inhibition occur multiple times in ∼every cell. The frequent direction reversals occurring after ∼2-10 minute intervals provided us the opportunity to compare forward versus reverse trajectories in the same oscillation as well as trajectories from different oscillations at different times during the movie.

Strikingly, both forward and reverse trajectories from the same oscillations and forward and reverse trajectories from different oscillations frequently retrace the same or similar curvilinear paths (**Fig. 2d, Movie 2**). This includes an example in which a return trajectory is different from its immediately preceding away from NS trajectory, but which instead overlaps with several preceding trajectories (**Fig. 2d**).

### Curvilinear long-range movements are dependent on the HSF1 transcription factor

Both the stable association with NS and the appearance of long-range curvilinear movements after transcriptional inhibition of the HSPA1 genes and transgenes were observed at 37°C and after heat shock (HS). However, at 37°C the HSPA1 genes express at ∼200-fold lower levels then when fully induced during HS and HSF1, the major transcription factor activating heat shock genes, exhibits much lower DNA binding to heat shock elements (HSEs) as compared to after heat shock ^31–33^.

To test whether the observed HSPA1 BAC transgene oscillations were dependent on the HSPA1 genes themselves, we generated deletion constructs removing the three HSPA1 genes ( 21.5kb deletion) plus the neighboring enhancer region (30.8kb deletion) from the original HSPA1 BAC transgene **(SFig. 3a**). Comparing several independently derived integrations of the wild-type (wt) or deletion HSPA1 BACs, we observed a decreased NS association frequency for the cell lines containing the deletion BAC transgenes **(SFig. 3b**). For each BAC stable integration (wt or one of the two deletions), we selected a cell clone that showed the transgene locating near the nuclear periphery in at least a fraction of cells (SFig. 3c&d); the logic here was to select clones with BAC integration genomic sites outside of NS-associated genomic regions. The wt BAC, as previously demonstrated^15,24^, targets near NS in all clones isolated, resulting in much lower frequencies of alleles mapping near the nuclear lamina.

**Figure 3.**
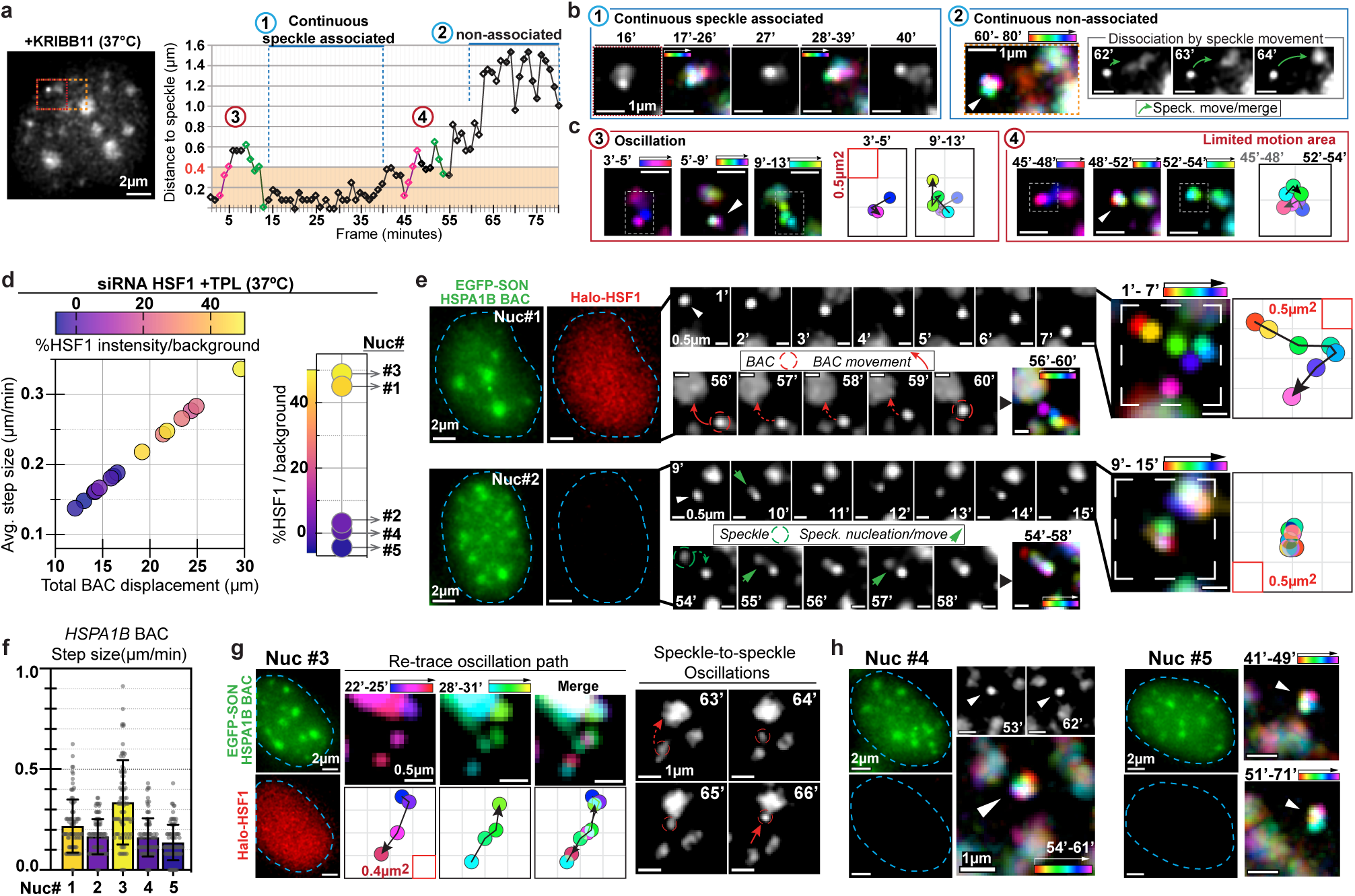
Curvilinear long-range *HSPA1B* BAC transgene movements are dependent on HSF1. **(a-c**) HSF1 inhibitor KRIBB11 treatment blocks *HSPA1B* transgene directional/curvilinear movements. (**a**) Time series of BAC-NS distance after KRIBB11 treatment, showing four apparent states: (1) steady BAC-NS association; (2) prolonged dissociation with restricted BAC movement; (3) apparent oscillation; (4) indeterminate/random movements. (**b)** Color-coded temporal projections showing state (1) steady NS-association and state (2) continuous non-association with NS (20 mins) and restricted transgene movement. During state (2) initiation (62-64 min), NS retraction rather than transgene movement drives separation. (**c**) Color-coded projections plus spatial grid of transgene position and trajectory show state (3) apparent oscillation with non-linear, more random transgene movements, and limited random movement during the state (4), contrasting with curvilinear movements in triptolide or α-amanitin-treated cells (see Fig. 2). Dashed boxes in projected images indicate gird area and are centered on transgene positions rather than NS. (**d-h**) HSF1 knockdown (KD) dose-dependently inhibitits long-range, curvilinear/directional *HSPA1B* transgene movements. (**d**) Left: Average frame-to-frame transgene step size (y-axis) versus total displacement (sum of all steps; x-axis); each bubble represents one nucleus (89 frames, 1 frame/min). Color indicates nuclear HSF1 expression level by fluorescence (% intensity above background; −6 to 50%) after KD. Right: HSF1 levels for labeled nuclei #1-5; nuclei # 1 and #3 show near-control HSF1 levels, while nuclei #2, #4, $5 show pronounced KD (see SFig. 4f). (**e**) Top: Long-range, curvilinear *HSPA1B* transgene trajectory in nucleus #1 with relatively high, uniform HaloTag-HSF1 fluorescence (red; left). Bottom: Near stationary *HSPA1B* transgene trajectories in nucleus #2 with low HSF1 fluorescence levels (left) (54-58 min shows NS movement with largely stationary transgene). (**f**) Transgene step size per nucleus. Bar height shows mean step size (μm) ±SD; scatter points represent individual step measurements (n = 88 per nucleus; nuclei #1–5). Nuclei with higher HSF1 levels (#1 and #3) display larger distribution range and increased frequency of larger step sizes compared to nuclei with lower HSF1 levels (#2, #4, #5). (**g-h**) Representative transgene trajectories from nucleus #3 (high HSF1) showing curvilinear movement including NS-to-NS linear movement (**g**), versus nuclei #4 and #5 (low HSF1) showing near-stationary transgenes (**h**).

Live-cell imaging (**SFig. 3e-i**) showed that cells carrying the wt BAC frequently oscillated in curvilinear motions in which the path moving away from the NS overlapped with the path moving back towards the NS **(SFig. 3e&h)**. In contrast, BACs with deleted *HSPA1* genes showed reduced oscillations (**SFig. 3h**), with the transgene instead frequently remaining peripheral and/or moving randomly **(SFig. 3f-g**). Instead, speckle attachment in these deletion BACs often occurred after their random collision with NS, movement of NS towards the transgene, or nucleation of NS adjacent to the BAC. Compared to the wt BAC, live-cell movies revealed these deletion BACs maintained significantly greater distances from speckles following triptolide treatment **(SFig. 3i, left panel)** and exhibited reduced mobility with smaller average step sizes **(SFig. 3i, right panel).** We concluded that both the NS targeting and oscillatory movements of the BAC transgenes were dependent on the presence of the HSPA1 genes.

Therefore, we next used both chemical inhibition and RNAi to test whether *HSPA1* gene oscillations were dependent on the acidic transcription factor, Heat Shock Factor 1 (HSF1). KRIBB11 was identified in a chemical screen for inhibitors of heat shock-induced luciferase expression driven by a HSE-containing promoter; subsequent analysis revealed that KRIBB11 inhibits heat shock induction of multiple heat shock genes and binds directly to HSF1 in vitro^34^. Binding of KRIBB11 is associated with reduced HSF1-mediated recruitment of p-TEFb to heat shock gene promoters for polymerase release, but KRIBB11 binding to HSF1 does not affect HSF1 trimerization or DNA binding.

Similarly to RNA pol 2 inhibition, KRIBB11 significantly decreased steady-state *HSPA1* NS association in fixed cells, as measured by the increased distances of both the endogenous locus and the *HSPA1B* BAC transgene from NS (**SFig. 1a-b**).

However, live-cell imaging revealed notable differences following treatment of cells with KRIBB11 versus RNA pol 2 inhibitors. BAC transgenes exhibited reduced movements throughout the movies and smaller step sizes compared to the control (**SFig. 4a, Movie 3**). Consequently, their total displacement per hour was lowest after KRIBB11 treatment. The sum of individual BAC steps within an oscillation (Δr) deviated from the expected ∼2 times the maximum NS separation distance expected for linear movements, consistent with a more random movement than that observed after RNA pol 2 inhibition (**Fig. 2c** vs **SFig 4b**). Additionally, as compared to after TPL treatment, after KRIBB11treatment the oscillation frequency decreased while the oscillation period increased (**SFig. 4c, Movie 3**). Because of the low BAC mobility and more random, nonlinear trajectories, “oscillations” were defined using changes in the BAC transgene-NS distance, without regard to whether the gene versus the NS was moving (**Fig. 3a, SFig. 4d**). Notably, whereas oscillation frequencies and durations were relatively constant among different nuclei treated with TPL, they varied more widely between nuclei after KRIBB11 treatment (**SFig. 4e**).

**Figure 4.**
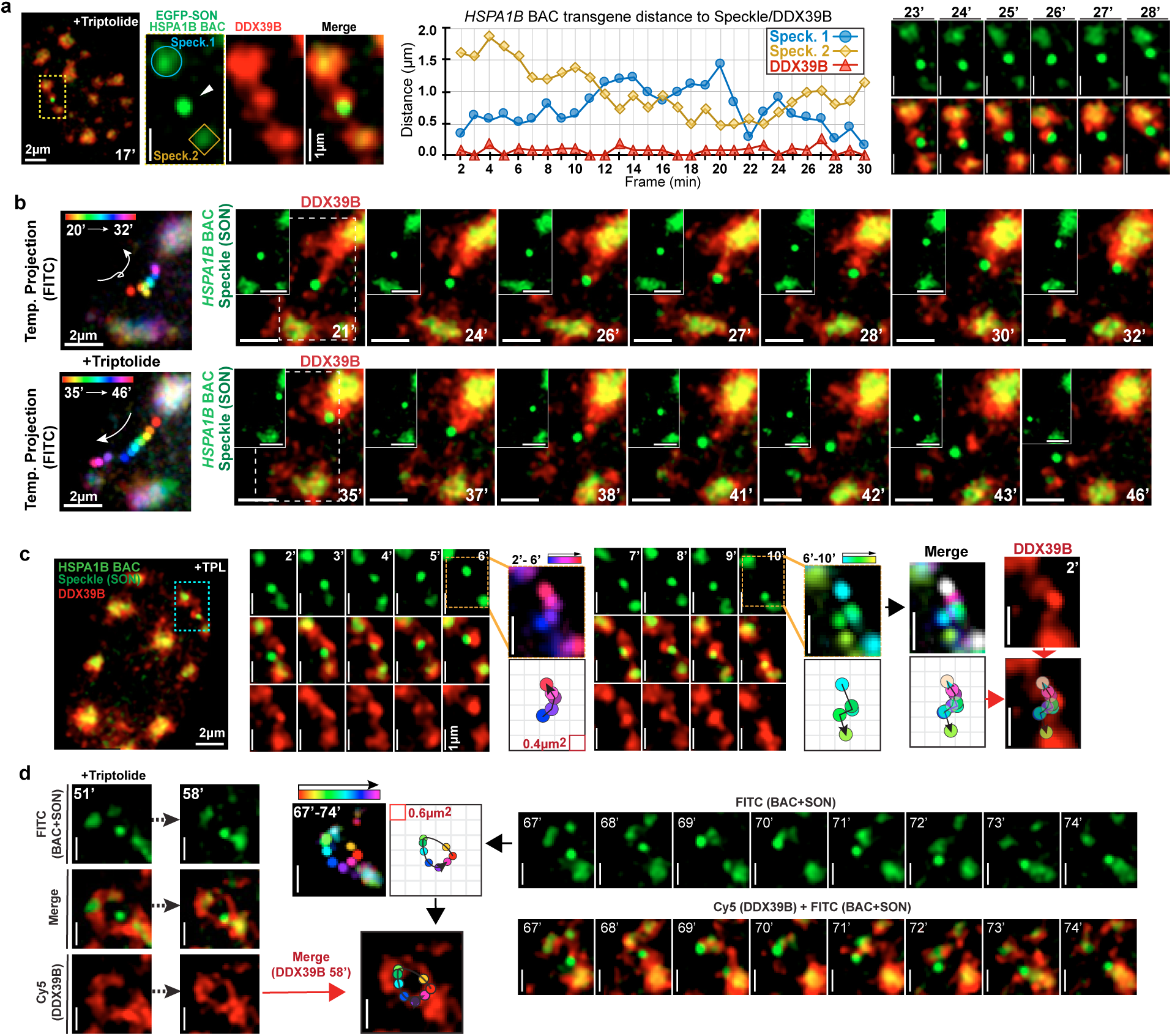
*HSPA1B* BAC transgene oscillations correlate spatially and temporally with DDX39B condensate dynamics. **(a-d)** Representative live-cell movies of four nuclei after triptolide treatment (37°C, 1 frame/min) showing HSPA1B BAC transgene (bright green), nuclear speckles (NS, green), and DDX39B (red). (**a**) Example (Movie 6) showing *HSPA1B* transgene (white arrow) in close association with DDX39B condensate during movement between two NS. Distance time series shows transgene in appearent constant contact with DDX39B, and movement towards one NS temporally coincides with DDX39B contraction towards the same NS. (**b**) Temporal color-coded projection of transgene revealing long-range curvilinear movements towards one NS (top) and away towards a smaller NS (bottom). Top: transgene movement towards one NS (21–32 min) correlates with DDX39B condensate retraction; second curvilinear movement away from that NS correlates with new DDX39B protrusion and foci movement away from the same NS (35–46 min). Scale bar = 2 μm. (**c**) *HSPA1B* transgene maintaining contact with DDX39B protrusion/foci during movements between two NS. Temporal color-coded projections and spatial grids show transgene trajectory: movement away from one NS and towards another (2-6 min), followed by return movement (6-10 min). Notably, transgene curved trajectories superimpose on the curved DDX39B connection between the two NS visualized at 2 mins (bottom right). (**d**) Example in which the temporal projection of a circular *HSPA1B* transgene curvilinear movement (67-74 mins) aligns with the circular path of DDX39B condensate connections between two NS visualized at earlier timepoint (58 mins). Scale bars = 1 μm.

In contrast to the curvilinear BAC movements, often with retraced oscillatory paths, induced after RNA pol 2 inhibition (**Fig. 2**), detailed analysis after KRIBB11 treatment revealed three distinct states of gene-NS association (**Fig. 3b&c**): (1) steady BAC association with NS resembling control; (2) prolonged dissociation marked by restricted and apparently random BAC transgene movement; and (3) “oscillations” resulting from a combination of random transgene movement, a relatively stationary transgene with curvilinear or random NS movement, or NS-initiated dissociation sometimes followed by BAC immobility.

To test the effect of HSF1 knockdown (KD) on *HSPA1B* transgene NS association and oscillatory movements, we fluorescently tagged endogenously expressed HSF1 using a CRISPR Halo tag knock-in strategy. Quantifying the nuclear HSF1 Halo tag intensity relative to the background, enabled direct measurement of relative KD efficiency in individual cells, both in fixed samples (**SFig. 4f**) and live cells prior to imaging (**Fig. 3 d&e)**.

HSF1 KD did not significantly change the HSA1B transgene distance distribution to NS, either in untreated cells or after TPL treatment. **(SFig. 4g**), as measured in fixed cells. In contrast, live-cell imaging after TPL treatment revealed a dose-dependent reduction in both average step size and total displacement of the *HSPA1B* transgene after HSF1 KD (**Fig. 3d)**.

Analysis of individual nuclei showed that cells with strongly reduced HSF1 levels after KD (**Fig. 3d right panel, Movie 4)** had largely stationary *HSPA1B* transgenes (**Fig. 3e&h**; nuclei #2 and #4-5), with NS association events frequently arising from NS movement and/or the nucleation of new NSs adjacent to the transgene (**Fig. 3e**, Nuc #2, green arrowheads). Quantitative measurements further confirmed that in cells with low HSF1 levels after KD, most BAC steps were restricted and <0.4 µm **(Fig. 3f)**. Conversely, nuclei retaining higher HSF1 levels after KD exhibited more mobile BAC loci with curvilinear oscillatory trajectories (**Fig. 3e&g**, nuc#1 and #3), and evidence of path retracing rather than randomly directed movements (**Fig. 3g**, nuc#3, middle panel).

These combined results from analysis of BAC transgenes with deletions of the HSPA1 genes, KRIBB11 treatment, and HSF1 KD lead us to conclude that long-range, curvilinear movement of HSPA1 gene loci is dependent on HSF1.

### Curvilinear *HSPA1B* transgene movements persist independent of actin, myosin VI, or cohesin perturbations

Previous work had suggested that disruption of actin polymerization and/or expression of a dominant negative actin targeted to the nucleus inhibited long-range, directional chromosome movement following VP16 transcriptional activation domain tethering, HS induction of HSPA1A plasmid transgenes, or induction of a U2 snRNA minigene tandem array^14,30,35^. A more recent study in budding yeast demonstrated that actin polymerization was required for INO1 gene repositioning upon INO1 activation^36^. However, these conclusions of an involvement of polymerized actin in directional chromosome movements were based on indirect inferences largely from fixed cell measurements rather than direct analysis of chromosome dynamics in live cells.

Here, we investigated the possible role of actin in the long-range *HSPA1B* transgene movements following transcriptional inhibition using both fixed and live-cell imaging. Additionally, both fixed and live-cell imaging suggested that nuclear myosin VI is involved in the clustering of RNA pol2 into distinct foci, possibly through long-range movements of genes^37^. Because RNA pol2 foci are concentrated surrounding nuclear speckles^38^, this suggested a possible involvement of nuclear myosin VI in gene movement relative to NS. Therefore, we also tested the possible role of nuclear myosin VI in the long-range *HSPA1B* transgene movement following transcriptional inhibition.

Actin polymerization was disrupted using either latrunculin A (LatA) or cytochalasin D (CytoD) treatment or by expression of a nonpolymerizable G13R actin mutant^39^ fused to mRFP and a nuclear localization sequence(NLS) (mRFP-NLS-tagged G13R β-actin)^30^. LatA binds actin monomers to prevent polymerization^40^ while CytoD caps the actin filament barbed end to block elongation. As controls for the dominant negative nonpolymerizable nuclear actin construct, we also expressed similar constructs containing either the wild-type β-actin sequence or a S14C actin mutant which enhanced F-actin stability and the F-actin/G-actin ratio *in vivo* ^39^. Expression of these nuclear actin constructs was validated via mRFP fluorescence in C7 CHO cells (C7MCP precursor lacking mCherry-MS2bp), while effective F-actin depolymerization after LatA or CytoD treatment was confirmed by phalloidin staining (**SFig. 5a**). Myosin VI function was inhibited using triiodophenoxy propionic acid (TIP), which blocks myosin VI ATPase activity and motor activity *in vitro* and *in vivo*^37,41^.

**Figure 5.**
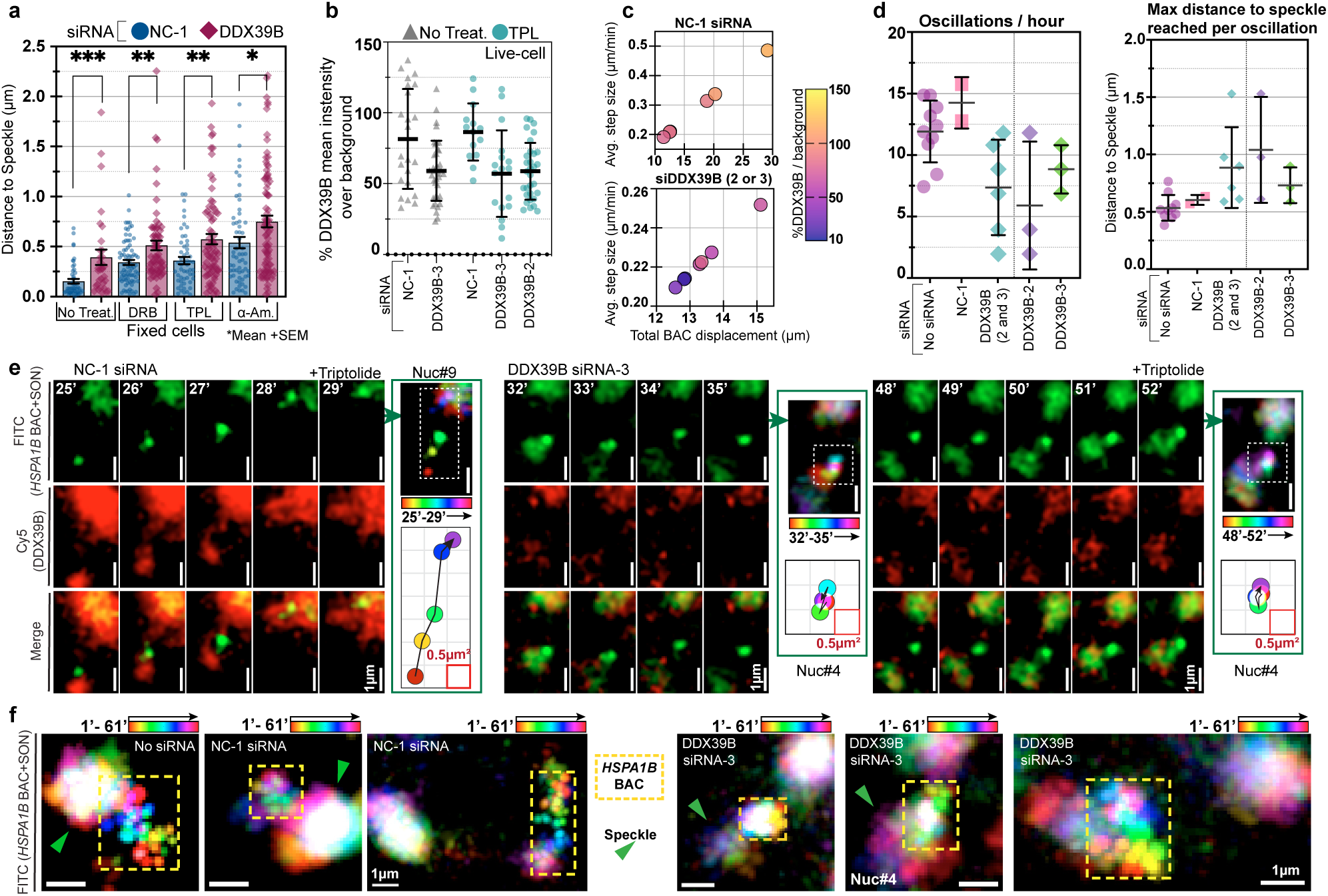
Increased *HSPA1B* distances from NS and dose-dependent reduction of *HSPA1B* curvilinear trajectories after DDX39B knockdown. (**a**) siRNA knockdown (KD) of DDX39B results in increased NS-*HSPA1B* transgene distance under control and inhibitor-treated conditions compared to NC-1 siRNA control. Mean distance to NS (μm) ±SEM with individual points measurements. Mann-Whitney U test; ns, P > 0.05; *P ≤ 0.05; **P ≤ 0.01; ***P ≤ 0.001; ****P ≤ 0.0001. NC-1 siRNA: n = 50, 76, 53, 60; DDX39B siRNA: n = 32, 65, 72, 85 (No Treat., DRB, TPL, α-amanitin, respectively). (**b**) DDX39B levels after siRNA KD in live cells measured as percentage of mean nuclear fluorescence intensity above background. Points represent individual nuclei; error bar shows mean ±SD. Large inter-nuclei variation of DDX39B levels in all conditions. Mean levels and range maintained across NC-1 and DDX39B siRNA with no treatment (left) and TPL treatment (right). Mean DDX39B reduction from NC-1 to DDX39B siRNA is consistent regardless of treatment.. (**c)** Reduced *HSPA1B* transgene net movements in cells with lower DDX39B levels after TPL. Bubble plot shows average step size (μm, y-axis) versus total displacement – sum of all steps (x-axis) – per nucleus (61 frames, 1 frame/min). Color indicates nuclear DDX39B level ( % above background; 10-150%). Top: NC-1 siRNA nuclei (n = 6). Bottom: DDX39B siRNA nuclei (n = 7; 3 with DDX39B-2, 4 with DDX39B-3 siRNA). (**d**) Reduced oscillation frequency with increased maximum BAC–NS distance after DDX39B KD. Left: Oscillations per hour per nucleus. Right: Mean BAC–NS distance per oscillation per nucleus. Both: no siRNA n = 10, NC-1 siRNA n = 2, DDX39B siRNA n = 6. Middle section shows combined DDX39B data; dotted line separates DDX39B-2 and DDX39B-3 (3 nuclei each). Bar height shows mean ±SD. (**e)** NC-1 siRNA control nucleus (left, Nuc #9) shows curvilinear transgene movements coordinated with DDX39B condensate dynamics after TPL, contrasting with reduced transgene mobility and random movements in DDX39B KD nucleus (middle and right, Nuc #4). Notably, apparent “oscillations” after DDX39B KD often occur due to NS movement away (32–35 min, middle) or towards (48–52 min, right) the relatively stationary transgene. (**f**) Color-coded temporal projections (1–61 min) comparing control (left; 1 no siRNA, 2 NC-1 siRNA) versus DDX39B-3 siRNA KD nuclei (right) after TPL. Control shows recurrent linear transgene excursions retracing similar spatial paths with static NS (white). DDX39B KD show restricted, less spatially loscalized transgene mobility with stationary BAC (white) and variable speckle positioning.

Fixed-cell analysis of NS-transgene distances revealed no significant changes after LatA, CytoD, or TIP treatment both in the absence or presence of TPL (P>0.05) (**SFig. 5b**). Similarly, no significant NS-transgene distance changes were observed following TPL treatment with expression of any of the S14C, G13R, or wild-type actin constructs (P>0.05) (**SFig. 5c**).

Live-cell imaging was unexpectedly complicated by an increased phototoxicity after combining any of the actin perturbations with TPL transcriptional inhibition, leading to a notable increase in the frequency of apoptosis. Despite these technical challenges, the characteristic long-range, oscillatory movements of the BAC transgene persisted across all experimental groups.

We observed oscillations and curvilinear transgene movements in S14C and G13R nuclear-targeted actin expressing cells similar to those in control cells, including between NS (**SFig. 5d**). However, expression of the S14C actin mutant produced noticeably increased NS mobility, accompanied by frequent NS fusion and fission events and an overall reduction in the number of NS per nucleus (**SFig. 5e**). This reduced number of NS, with a concomitant increase in distances between NS, may explain the observation of unusually long curvilinear transgene movements between NS observed in some cells expressing the S14C actin mutant (**SFig. 5d right & 5e**). Similarly, live-cell imaging of DRB-treated cells demonstrated continued transgene oscillations and curvilinear trajectories after LatA treatment (**SFig. 5f**). Additionally, we compared BAC transgene oscillation dynamics with either TPL treatment alone or additional treatment with either CytoD, or TIP (n=3 per treatment). No significant differences in mean NS-transgene distance or transgene step size (μm/min) were observed between these conditions (P > 0.05), while the oscillation frequency remained similar across treatments as well as the maximum transgene-NS separation distances and total transgene displacement per oscillation (P > 0.05) (**SFig. 6a-b**).

Recently, cohesin has been suggested as contributing to the tethering of genes to NS^18^. Given the known loop-extrusion activity of cohesin, we therefore next asked whether cohesin might drive *HSPA1B* BAC transgene oscillations. We used CRISPR insertion of the HaloTag to tag the endogenously expressed RAD21 protein, allowing us to assess the extent of RAD21 siRNA KD in both fixed and live cells **(SFig. 6c left).** We observed no significant change in NS-transgene distances in fixed cells following RAD21 KD and then TPL treatment (P > 0.05) (**SFig. 6c right**). Live-cell imaging did reveal altered nuclear dynamics, including frequent nuclear constriction and expansion, possibly in cells undergoing mitotic slippage^42^, together with high NS motility, NS fusion, and reduced NS numbers **(Movie 5)**. Despite these dramatic nuclear architectural changes, we observed typical curvilinear, long-range *HSPA1B* transgene movements even in cells with substantial RAD21 KD **(SFig. 6d-e, Movie 5).** Quantitative analysis (n = 3 nuclei) showed similar oscillation frequency per hour and no significant changes in various parameters characterizing these oscillation dynamics (P > 0.05) (**SFig. 6d**).

We therefore conclude that the long-range, curvilinear *HSPA1B* oscillatory movements are independent of F-actin, myosin VI, and/or cohesin based mechanisms.

### Curvilinear *HSPA1B* transgene movements spatially and temporally correlate with DDX39B condensate dynamics

Perhaps the most intriguing aspect of the long-range oscillatory movements of *HSPA1* genes after transcriptional inhibition are the extended curvilinear trajectories between adjacent NS, especially pronounced for the *HSPA1B* BAC transgene. These trajectories spanned nearly the entire nuclear width in some cells expressing the S14C actin-NLS mutant construct in which the NS number was reduced, resulting in widely spaced NS **(SFig. 5e**). In a parallel study, we discovered DDX39B, together with several Exon Junction Complex (EJC) related proteins, formed highly dynamic concentrations extending outwards from NS and often bridging subsets of adjacent NS^19^. While the exact physical nature of these DDX39B concentrations outside NS remains to be determined, as they do not show the expected rounded shape predicted for liquid-liquid phase separation, for clarity we will refer to these structures as DDX39B condensates. Additionally, we observed that highly expressed chromosomal regions, including those with strong or weak NS attachment, frequently associated with these DDX39B condensates^19^. In a second parallel study, we showed by live-cell imaging that small SON condensates translocate between adjacent NS within these bridging DDX39B protein condensates^43^.

Therefore, we next examined the spatial and temporal correlations between this DDX39B condensate network and curvilinear *HSPA1B* transgene trajectories between NS. We first used CRISPR to tag the endogenously expressed DDX39B protein with HaloTag in the CHO C7MCP cell line (**Fig. 4a**). Live-cell imaging after TPL treatment demonstrated that oscillation dynamics and curvilinear transgene trajectories were preserved, with continuous oscillations observed in 42 out of 43 nuclei.

Notably, after dissociation from NS, the *HSPA1B BAC* transgene was typically associated with a DDX39B condensate—even if only as a small, isolated droplet adjacent to the transgene (37 out of 43 nuclei, **Fig. 4a, Movie 6**). Strikingly, *HSPA1B* transgene movements between NS connected by DDX39B condensates exhibited both spatial and temporal correlation with DDX39B dynamics (**Fig. 4a&b, Movie 6**). Specifically, transgene movement away from one NS and towards another consistently aligned with protrusions or movements of the DDX39B condensate in the same direction (**Fig. 4b**, bottom panels, **Movie 6**). Similarly, return movements of the *HSPA1B* transgene towards a NS correlated with retractions of the DDX39B condensate back into the NS (**Fig. 4b**, top panels, **Movie 6**). Furthermore, periods of transgene immobility often coincided with it being connected by an isolated small DDX39B droplet rather than a DDX39B extension connected to a nearby NS (**SFig. 7a**).

Additionally, the curvilinear paths of transgene oscillations visualized by projections over time closely matched the curvilinear path connecting multiple DDX39B accumulations, as visualized even at a single time point; thus, the transgene trajectory path aligned with the shape of the pre-existing DDX39B connection at a given time (**Fig. 4c**). DDX39B condensates consistently occupied and appeared to move along specific routes over time (**Fig. 4d**). Thus, the dynamics of DDX39B condensates bridging between NS mark appear to mark pre-determined paths for the HSPA1B transgene movements.

In summary, these results show first that *HSPA1B* transgenes locate immediately adjacent to DDX39B condensates and second that the curvilinear movements of *HSPA1B* transgenes both spatially and temporally tightly correlate with the dynamic extensions, retractions, and translocations of these DDX39B condensates.

### Curvilinear *HSPA1B* transgene oscillation dynamics are dependent on DDX39B levels

Given the extensive spatial and temporal correlation between *HSPA1B* transgene and DDX39B condensate dynamics, we next asked whether *HSPA1B* transgene curvilinear movements and oscillations were dependent on DDX39B expression.

Cells were transfected with either a non-targeting siRNA negative control (NC-1), or one of two DDX39B-targeting siRNAs (#2 or #3), which both produced similar knockdown efficiency (data not shown). In fixed cells, DDX39B knockdown (KD)—but not the NC-1 siRNA control—significantly increased the distances between the *HSPA1B* transgene and NS in untreated samples as well as in those exposed to each of the three RNA pol 2 inhibitors (**Fig. 5a**).

Live-cell imaging showed variable DDX39B signal intensities per cell above background but with a mean decrease in the overall DDX39B (rather than DDX39B condensate) per cell of ∼1.4-fold for both of the DDX39B siRNAs compared to the NC-1 controls (**Fig. 5b**). We analyzed live-cell imaging data only from cells retaining moderate levels of DDX39B that retained stable SON-marked NS, because in cells with lower levels of DDX39B we frequently observed SON-marked NS partially dispersing, disappearing, and reappearing over several 1-minute intervals (**SFig. 7b, Movie 7**). Based on fixed cell images, we suspect this phenotype is followed by major reorganization of chromatin, dispersion of SON to the interchromatin space, and then apoptotic cell death (**SFig. 7c**).

Thus, we selected nuclei with a DDX39B intensity over background of 10–80% for DDX39B siRNA-transfected cells, and 70–115% for NC-1 controls. Following TPL treatment, transgene mobility showed a DDX39B dose-dependent decrease with smaller average step sizes and total BAC displacement correlating with lower DDX39B levels (**Fig. 5c**). In contrast, control (NC-1) siRNA treated nuclei showed increased average step sizes and total BAC displacement in cells expressing higher DDX39B levels (**Fig. 5c, top**). Additionally, NC1-treated nuclei displayed a higher oscillation frequency, similar to non-transfected controls, while DDX39B siRNAs produced fewer oscillations (**Fig. 5d**, left panel). The average distance from NS reached within each oscillation was similar for no-siRNA and NC1 nuclei, but increased in DDX39B siRNA-transfected nuclei (**Fig. 5d**), consistent with fixed cell measurements (**Fig. 5a**).

Closer examination revealed *HSPA1B* transgene movements appeared largely random after DDX39B KD. The apparent “oscillatory” behavior-derived from plots of NS distance over time-instead appeared to be primarily the result of NS movements and/or NS nucleation events rather than directed, curvilinear transgene trajectories. Throughout these “oscillation” intervals, the transgene exhibited limited or no contact with residual DDX39B condensates (**Fig. 5e**, right two panels) in contrast to the curvilinear movements observed after the control NC-1 siRNA KD (**Fig, 5e**, left panel). Projecting images captured every minute for 60 minutes into a single frame in control cells showed relatively stable NS positions (white) adjacent to elongated tracks containing the multi-colored transgene locations, consistent with their repeated curvilinear oscillatory movements along the same or similar paths (**Fig. 5f**, left panel. In contrast, in DDX39B KD cells we observed white signals-reporting locations where the transgene was relatively stable-alongside variable colored areas indicative of more erratic NS movement (**Fig. 5f**, right panel).

Therefore, we conclude that the long-range, oscillatory, curvilinear movements of the *HSPA1B* transgenes are dependent on DDX39B expression and DDX39B condensate dynamics.

## Discussion

Here we set out to understand the stable association of the HSPA1 locus with NS. Live-cell imaging revealed first that the reduction in NS association of HSPA1 genes and transgenes observed after RNA pol 2 inhibition was due to loss of stable NS binding. Second, live-cell imaging revealed a constitutive mechanism for long-range, directional HSPA1 gene movement to and from NS dependent on the HSF1 transcription factor that activates HSPA1. These results suggest a model in which stable NS association of the HSPA1 gene locus and transgenes is produced by the combined action of two distinct molecular mechanisms-one conferring prolonged binding to NS and another conferring an active, directional long-range gene movement to return HSPA1 genes back towards NS whenever they become detached. Because the HSPA1 prolonged binding to NS is reduced after treatment with three different RNA pol 2 inhibitors, the simplest model would be that nascent RNA transcripts are required for the prolonged NS binding activity of HSPA1 genes. These nascent RNA transcripts might be produced by transcription from flanking house-keeping genes at normal temperatures but by HSPA1 transcripts after heat shock.

Whereas previously F-actin and nuclear myosins had been implicated in similar long-range, directional gene movements, here we used live-cell imaging to rule out a requirement for F-actin, nuclear myosin VI, as well as cohesin. Instead, our results suggest that it is the dynamics of DDX39B condensates, extending and retracting from NS, which drive this HSPA1gene movement. The temporal and spatial correlation of HSPA1 gene movement with DDX39B condensate dynamics provides an explanation for the unexpected observation that HSPA1 gene movements towards and away from NS frequently retrace the the same or similar curvilinear path during the repeated HSPA1 gene oscillations observed after transcriptional inhibition.

By manipulating the growth and shrinkage of artificial condensates designed to bind chromatin, a recent study demonstrated the ability of capillary forces generated by condensate dynamics to drive chromatin movements^44^. Here we are now tying a physiologically important response-the efficient HS induction of *HSPA1* genes - to long-range gene movements driven by endogenous condensate dynamics. We previously demonstrated a multi-fold gene expression amplification of *HSPA1* gene heat shock induction that is dependent on contact with NS^14,15^. By providing a mechanism for rapidly restoring *HSPA1* NS association within several minutes for the small percentage of *HSPA1* alleles that lose NS attachment, these HSPA1 long-range gene movements provide an important robustness to the HS response that is directly tied to cell survival.

There are several current limitations of this study. So far, we only examined the NS association and long-range gene movement for the *HSPA1* gene locus and *HSPA1* BAC transgenes, with much of our analysis focused specifically on the BAC transgene system. We do not know yet the exact physical nature of what we are calling the DDX39B condensate; moreover, the properties of this condensate may be dependent on multiple proteins, with these dependences possibly varying in different cell types and lines. However, current strengths of this study are that we have demonstrated the existence of an active, transcription-factor dependent, and nuclear condensate dependent mechanism that drives long-range gene motion over micron-scale distances and several minute time intervals.

For several reasons, we suspect that our observations of transcription-factor mediated, DDX39B condensate driven gene motion are likely to extend to other genes and chromosome regions. We previously described similar long-range, directional chromosome movements after tethering of the VP16 acidic activation domain (AAD)^30^. HSF1 is an acidic transcription factor with a strong AAD. In a parallel study, we recently demonstrated that NS-targeting activity is a common activity of many AADs that is reduced after transcriptional inhibition^45^. In a second parallel study, we showed the colocalization of highly expressed chromosome regions adjacent to DDX39B condensates and NS^19^.

We therefore anticipate future investigations to further explore the role of DDX39B condensate dynamics may play in nuclear genome organization and specifically in gene positoning and chromosome motion.

## Materials and Methods

### Cell Culture

HCT116 and CHO-K1 cells were maintained in ATCC McCoy’s 5A or Ham’s F12 medium, respectively (Cell Media Facility, University of Illinois Urbana-Champaign), supplemented with 10% (v/v) fetal bovine serum (FBS; Sigma-Aldrich, F0926) at 37°C in a humidified incubator with 5% CO2. Cells were passaged at ∼80% confluency by washing once with phosphate-buffered saline (PBS), then dissociating with 0.05% Trypsin-EDTA (Gibco, 25300062) for HCT116 or 0.25% Trypsin-EDTA (Gibco, 25200072) for CHO-K1. Trypsin was inactivated with complete medium and cells were split as needed.

The **CHO-K1 C7MCP** cell line was generated as previously described by sequential transfection with the *HSPA1B*-3’UTR-MS2 BAC, p3’SS-EGFP-dLacI, the EGFP-SON BAC, and then the pUB-MS2bp-mCherry plasmid ^15^. Selection was maintained with 200 µg/mL G418 (Gibco, Cat.# 10131035), 200 µg/mL Hygromycin B (Thermo Fisher Scientific, Cat# 10687010), and 100 µg/mL Zeocin (Gibco, R25001).

The **HCT116 96-8** cell line was generated as previously described^27^; this line contains a 96-mer TetO repeat integrated via CRISPR/Cas9 knock-in ∼16 kb downstream of the endogenous *HSPA1B* gene. Then CRISPR-mediated knock-in was used to endogenously tag SON protein at the N-terminus; single-cell clones were isolated to generate the HCT116 96-8 B2 cell line. The pF9-TetR-Halo-NLS plasmid was stably transfected into HCT116 96-8 B2 cell line and selected with 200 μg/mL hygromycin B for 2 weeks.

### CRISPR/Cas9 Knock-in

A Cas9/sgRNA ribonucleoprotein (RNP) complex and linear dsDNA donor were transfected using the Amaxa Nucleofector II device (Lonza) with the Cell Line Nucleofector Kit V (Lonza, Cat# VCA-1003) and program D-032 for HCT116 cells, or Kit T (Lonza, Cat# VCA-1002) and program U-023 for CHO-K1 cells. For each transfection, 0.5 million cells were pelleted and resuspended in 90 μL of nucleofection reaction buffer. In parallel, 3 μg of Cas9 protein (Alt-R S.p. Cas9 Nuclease V3, IDT, Cat# 1081058) and 2 μg of purified sgRNA were incubated in 10 μL of reaction buffer for 10–15 minutes at room temperature. The RNP complex was then combined with 5 μg of purified donor DNA and the 90 μL cell suspension, transferred to a nucleofection cuvette, and electroporated using the appropriate program. Cells were recovered in a 6-well plate with complete media, and the medium was replaced 12–24 hours post-transfection.

### sgRNA Generation

sgRNAs were generated by in vitro transcription (IVT) using a DNA template created via two-step overlap PCR. The first step assembled the sgRNA/tracrRNA fusion using a forward primer containing a T7 promoter sequence followed by a target-specific 20 bp sequence upstream of the PAM, and a 15bp overlap with tracrRNA (5’–TAATACGACTCACTATAGNNNNNNNNNNNNNNNNNNNNGTTTTAGAGCTAGAA–3’, where N represents the target-specific sgRNA sequence; see Extended Data Table 1 for target sequences) and a reverse primer containing the tracrRNA sequence (5’–AAAAGCACCGACTCGGTGCCACTTTTTCAAGTTGATAACGGACTAGCCTTATTTTAA CTTGCTATTTCTAGCTCTAAAAC–3’). The second step amplified the sgRNA template using a T7 promoter-containing forward primer (5’–TAATACGACTCACTATAG–3’) and a universal reverse primer (5’–AAAAGCACCGACTCG–3’) complementary to the tracrRNA sequence. The sgRNA DNA template was purified using the NucleoSpin Gel and PCR Clean-up Mini kit (Macherey-Nagel, Cat# 740609) and transcribed using in-house purified T7 polymerase. Post-IVT sgRNA was purified using the QIAGEN RNeasy Mini Kit (Qiagen, Cat# 74104) with modifications for short-length (<200 bp) RNA. After purification, sgRNAs were denatured at 95 °C for 5 minutes, cooled to 25 °C at 0.1 °C/s, and stored at −80 °C.

### Donor DNA Preparation

Donor DNA was amplified by PCR using Q5 High-Fidelity DNA Polymerase (New England Biolabs, Cat# M0491S) with modified 5’ amine-labeled primers (C6 linker^46^). Primers contained ∼60 bp homology arms flanking the target knock-in site immediately upstream or downstream of the start codon (ATG), plus sequence encoding a Halo-Tag and GSG linker; specific primer sequences are listed in Extended Data Table 1. For the HCT116 96-8 B2 line, donor DNA was amplified from a modified SON-containing BAC (RP11-165J2, Invitrogen) with EGFP fused to the N-terminus of SON^14^ using forward primer 5’–GTGCTCACTGATTGGTCCCTC–3’ and reverse primer 5’–GAACGACTGCGTCTCCGAAG–3’. Donor DNA was purified sequentially using the NucleoSpin Gel and PCR Clean-up Mini kit and NucleoSpin Gel and PCR Clean-up XS (Macherey-Nagel, Cat# 740611) to achieve a final concentration of ∼1 μg/μL.

### Single-Cell Cloning

Five days post-transfection, EGFP- or HaloTag-positive cells were sorted using a FACS Aria II sorter (Roy J. Carver Biotechnology Center, University of Illinois Urbana-Champaign) with single cells deposited into individual wells of a 96-well plate for single-clone cell line generation.

### Recombineering of HSPA1 BACs

#### Retrofitting of HSPA1 BAC vector (RP11-92G8-RFTD)

The vector backbone of a 180kb human HSPA1 BAC (RP11-92G8) was retrofitted via lambda Red-mediated recombination using previously published protocols^47^. A 10 kb retrofitting cassette was obtained from the RFTD-vector plasmid^45^ by PacI restriction digestion, purified by agarose gel extraction, and contained a 96-mer Tet operator (TetO) repeat, puromycin resistance marker, and kanamycin resistance marker. This cassette was flanked by 415 bp (5’) and 411 bp (3’) homology arms designed to target the BAC vector backbone for recombination (Extended Data Table 1). The E. coli strain SW102 was used, and recombinants containing the retrofitted vector were selected for kanamycin (100ug/ml) resistance in LB agar plates at 30 ^0^C. BAC constructs were then screened by restriction enzyme fingerprinting to verify BAC integrity and the absence of DNA deletions between repetitive elements generated during the recombineering process.

#### Generating HSPA1 Deletion BACs

We used two-step galK-based BAC recombineering to introduce deletions into the retrofitted RP11-92G8-RFTD BAC. The galK selection cassette was amplified by PCR from the plasmid pUGG template with 70–80 bp homology arms added through incorporation into the PCR primers; these homology regions corresponded to DNA sequences immediately upstream and downstream of the target deletion site (Extended Data Table 1). In the first step, the galK cassette was introduced into the BAC through homologous recombination, replacing the genomic region to be deleted, Positive selection was through growth on minimal medium plates containing galactose (0.2% w/v) as the sole carbon source. In the second step, the galK cassette was selectively deleted by homologous recombination using a DNA fragment containing seamlessly joined homology arms without the galK sequence and using negative selection by growth at 30 °C on minimal medium containing 2-deoxy-galactose (0.2% w/v), which is toxic to galK-expressing cells, and glycerol (0.2% v/v) as a carbon source. BACs from surviving colonies were screened by restriction enzyme fingerprinting and PCR verification of galK cassette absence. This approach generated the final deletion BACs RP11-92G8-RFTD-D3 and RP11-92G8-RFTD-D3B (also referred to as 92G8-RFTD-HSseD).

### BAC integration and clone selection of HSPA1 BAC containing cell lines

To generate additional CHO cell lines with HSPA1 BAC integrations, we started with the previously described CHO-K1 EGFP-SON-D6 cell line^48^. This line was created by transfection with a retrofitted human SON BAC (RP11-165J2, Invitrogen) containing an N-terminal EGFP tag and Zeocin resistance cassette, selecting for stably expressing clones. From this point forward, this line is referred to as CHO D6.

We next transfected CHO D6 cells with one of each of the three HSPA1 BAC constructs. The original BAC (RP11-92G8) was retrofitted to include a backbone cassette containing a 96-mer Tet operator repeat and a puromycin resistance marker^45^ creating the construct designated RP11-92G8-RFTD (referred to as wild-type BAC). This BAC was further engineered by recombineering to generate two deletion derivatives: 92G8-RFTD-D3 (Deletion of a 21.5 kb region encompassing all three HSPA1 genes; *HSP*Δ in figure) and 92G8-RFTD-HSseD (Deletion of a 30.8 kb region, which includes the first deletion as well as an additional downstream super-enhancer region identified near *HSPA1B; HSP*+EnhΔ in figure). CHO D6 cells were transfected with each BAC variant and selected using puromycin at 7.5 µg/mL (Gibco, Cat# A1113803). Following selection, subclones were isolated and cell lines established for each BAC construct derived from specific cell clones.

To screen these cell lines from specific cell clones, cells were transiently transfected with a pF9-TetR-Halo-NLS plasmid. Cells were labeled with a HaloTag ligand, and fluorescent TetO foci were evaluated by microscopy. Cell lines derived from specific clones were screened for diffraction-limited size spots (0.5 µm diameter), indicative of low copy number integrations, and preferential positioning near the nuclear lamina and thus distant away from nuclear speckles under control conditions at 37 °C. Our rationale was this would avoid cell clones in which the BAC had integrated into a nuclear speckle associated chromosome domain. Using this screening, we selected three specific cell lines: CHO D6 92G8-FULL-C1 (RP11-92G8-RFTD), CHO D6 92G8-D3-a1C (92G8-RFTD-D3), and CHO D6 92G8-HSseD-T (92G8-RFTD-HSseD). All of these HSPA1 transgene containing cell lines were maintained in medium supplemented with 5 µg/mL puromycin and 100 µg/mL zeocin to retain EGFP-SON expression and transgene integrations.

### DNA Transfection

BAC DNA for transfection was prepared using the QIAGEN Large Construct Kit (Qiagen, Cat# 12462) according to the manufacturer’s instructions. All other plasmid constructs were purified using either the NucleoSpin Plasmid Mini kit (Macherey-Nagel, Cat# 740588) or the NucleoBond Xtra Midi kit (Macherey-Nagel, Cat# 740410). For transfections, cells were seeded in T-75 cell culture flasks and grown to approximately 70% confluence. Lipofectamine 2000 (Thermo Fisher Scientific, Cat# 11668019) was used in serum-free media according to the manufacturer’s instructions, employing a 1:2.5 DNA-to-Lipofectamine ratio. A total of 10 μg purified BAC DNA or 7 μg plasmid DNA was used per transfection. For generation of stable integration cell lines, after 24 hours, the culture medium was replaced with fresh medium supplemented with appropriate drug selection. For transient transfections, cells were analyzed approximately 48 hours post-transfection without antibiotic selection.

### Mapping *HSPA1B* BAC integration in CHO C7MCP cells

To obtain high-molecular-weight (HMW) genomic DNA (gDNA) for long-read sequencing, C7MCP cells were grown to approximately 75% confluency. Following trypsinization and resuspension in culture medium, 1 × 10 cells were collected in a 2 mL tube, centrifuged, and resuspended in 200 µL of 1× PBS. HMW gDNA was extracted using the MagAttract HMW DNA Kit (Qiagen, 67563) according to the manufacturer’s instructions, yielding DNA fragments in the 10–50 kb range suitable for long-read sequencing applications. DNA concentration was determined by NanoDrop spectrophotometry (28.3 ng/µL) and Qubit fluorometry (22.8 ng/µL). DNA purity was assessed by NanoDrop (A260/280 = 2.17, A260/230 = 2.23), indicating high-quality DNA. A total of 2.2 µg of gDNA was submitted for PacBio long-read sequencing at the University of Illinois Roy J. Carver Biotechnology Center, DNA Services Laboratory for PacBio sequencing. The gDNA was sheared with a Megaruptor 3 to an average fragment length of 15 kb. Sheared gDNA was converted to a library using the SMRTBell Prep kit 3.0. Circular consensus sequencing (CCS) analysis was performed using SMRTLink v11.0 with the following parameters: *ccc - - m n - pcc c 3 -- m n - q 0.99*. Sequencing yielded a total of 2,375,131 reads with a mean read length of 17,042 bp.

#### Read alignment and BAC integration mapping

We identified the HSPA1B BAC integration site in CHO K1 C7MCP cells using PacBio HiFi sequencing and a sequence of custom bioinformatic analyses developed in collaboration with the High-Performance Computing in Biology (HPCBio) group at the Roy J. Carver Biotechnology Center, University of Illinois at Urbana-Champaign. This overall analysis was designed to detect chimeric reads spanning the junction between the HSPA1B BAC vector and the Chinese hamster genome.

Reads were aligned using Minimap2 to a hybrid reference genome combining the Chinese hamster genome assembly (*Cricetulus griseus*, CriGri-PICRH-1.0, NCBI Assembly GCF_003668045.3) with the HSPA1B and EGFP-SON BAC sequences. The EGFP-SON BAC was included to prevent ambiguous mapping due to shared backbone homology.

### Identification of Chimeric Junction Reads

Using Samtools, HiFi reads possessing a supplementary alignment (SA) tag were retrieved. We filtered for chimeric reads where the SA corresponded to the HSPA1B BAC and the primary alignment mapped to a hamster scaffold. The alignments were sorted and indexed for visualization in the Integrative Genomics Viewer (IGV). Candidate reads were grouped by their hamster scaffold position to identify clusters of chimeric reads, and candidate HiFi reads mapping to these distinct genomic loci were extracted for detailed sequence analysis.

#### Integration Site Verification and Mapping

Four candidate HiFi reads (ranging from15–22 kb in length) were independently identified as spanning the integration junction. These reads were analyzed via BLASTN against the standard Chinese hamster genome and the HSPA1B BAC construct using the NCBI BLAST 2 sequences web service. A split alignment profile—comprising 5–10 kb of hamster sequence flanking 5–10 kb of BAC sequence—identified the integration site at Chinese hamster chromosome X (NC_048604.1): 120,042,677–120,060,113.

#### BAC Integration Junction Characterization

To search for the second junction, we queried the HiFi dataset with the 5 kb sequence downstream of the identified insertion site. Retrieved reads were then BLAST-aligned back to the hamster genome assembly. However, all retrieved reads matched the wild-type hamster genome completely and lacked BAC sequences; consequently, we were unable to identify the second junction of the BAC integration.

### TSA-seq Mapping

TSA-seq (Tyramide Signal Amplification sequencing) was performed as described previously^16^. For the HCT116 cell line, TSA-seq tracks were obtained from published datasets^16^. TSA-seq for CHO-K1 cells was performed in-house.

### Drug Treatments and Heat Shock

For RNA pol 2 transcriptional inhibition, cells were treated either with 50 µg/mL DRB (5,6-dichloro-1-β-D-ribofuranosylbenzimidazole; Sigma-Aldrich, D1916), 0.1 µg/mL Triptolide (Sigma-Aldrich, 67563), or 50 µg/mL α-amanitin (Sigma-Aldrich, A2263). For inhibition of heat shock genes via inhibition of HSF1, cells were treated with 0.05 mM KRIBB11 (Sigma-Aldrich, 385570). For both fixed and live-cell samples, 50% of culture medium was removed, inhibitors were added at 2× final concentration to the removed medium, and the medium was returned to the dish. Stock concentrations were 50 mg/mL DRB, 1 mg/mL triptolide, and 20 mM KRIBB11 in DMSO, and 1 mg/mL α-amanitin in H O. For fixed cell experiments, cells were treated with DRB, Triptolide, or KRIB11 for 2 hours or α-amanitin for 5 hours prior to fixation. For live-cell microscopy experiments, cells were treated with DRB, Triptolide, or KRIBB11 for 60-90 minutes or α-amanitin for ∼3 hours, followed by approximatley 30 min for field selection and focal plane adjustment before initiating live-cell image aquisition.

For actin and myosin VI inhibition, cells were treated with 1 µM latrunculin A (Cayman Chemical, 10010630), 5 µg/mL cytochalasin D (Source, Cat.No.), or 25 µM 2,4,6-triiodophenol (TIP, myosin VI inhibitor; Source, Cat.No.), following the same medium addition procedure. Stock concentrations were 250 µM latrunculin A, 1 mg/mL cytochalasin D, and 250 mM TIP, respectively. Cells were treated with latrunculin A, cytochalasin D, or TIP for 1 hour prior to fixation or live cell imaging.

For heat shock treatment for fixed cell microscopy, cells were seeded on coverslips in 24-well plates or 35-mm dishes and grown to approximately 70% confluency. Dishes were sealed with parafilm and submerged in a water bath preheated to 42.6°C for 30 min without medium change. Cells were then immediately fixed with 2.0% paraformaldehyde (PFA) for 15 min and washed three times with 1× PBS for 5 min each before mounting coverslips with DAPI-DABCO medium.

### HaloTag Ligand Labeling

For labeling of Halo-tagged proteins, Janelia Fluor 646 HaloTag Ligand (Promega, GA112) was resuspended in DMSO to a stock concentration of 200 µM. For fixed-cell samples, the ligand was added at a 1:10,000 dilution (20 nM final concentration) and incubated for 2 hours, or at a 1:40,000 dilution (5 nM final concentration) and incubated overnight. For live-cell imaging, the ligand was added at a 1:5,000 dilution (40 nM final concentration) and incubated for a minimum of 1 hour before imaging. In both cases, labeled cells were returned to the incubator at 37°C before fixation or imaging.

### Phalloidin Staining

Cells were fixed with 2% paraformaldehyde (PFA) for 15 min at room temperature, then washed three times with 1× PBS for 5 min each. Cells were permeabilized with 0.1% Triton X-100 (Thermo Fisher Scientific, 28314) for 15 min, followed by three washes with 1× PBS for 5 min each. Cells were then incubated with Phalloidin-iFluor 647 (abcam, ab176759) at a 1:4,000 dilution and 1% bovine serum albumin (BSA; Source, Cat.No.) for 3 hours at room temperature. Following staining, cells were washed three times with 1× PBS for 5 min each, then mounted on coverslips with DAPI-DABCO mounting medium.

### RNA Interference-Mediated Knockdown

For gene knockdown via RNA interference, custom Dicer-substrate siRNA (DsiRNA) sequences (2 nmol, IDT) targeting individual genes and a non-targeting DsiRNA negative control (1 nmol, IDT, 51-01-14-03) were used. DsiRNAs were resuspended in Nuclease-Free Duplex Buffer (IDT, 11-01-03-01) to a final concentration of 20 µM, then heated to 94°C for 2 min before being allowed to cool to room temperature. Reconstituted DsiRNA solutions were stored at −20°C for future use. For siRNA sequences see Extended Data Table 1.

For fixed-cell experiments, cells were seeded into 24-well plates containing coverslips. For live-cell experiments, cells were seeded in equivalent volumes directly into 35 mm dishes with glass coverslip bottoms (MatTek). After 1–2 days of culture at approximately 40% confluency, cells were transfected with DsiRNA (20–30 nM final concentration) complexed with Lipofectamine RNAiMAX (Thermo Fisher Scientific, 13778030) at a 1:600 µL ratio in Opti-MEM Reduced Serum Medium. DsiRNA-Lipofectamine RNAiMAX complexes were incubated at room temperature for 20 min, then added in reduced volume (600 µL) to coat the coverslip, using the effective well created by the thickness of the plastic dish with the glass coverslip glued to its bottom surface. Culture medium was replaced 12–24 hours after transfection, and cells were incubated for a total of 24–48 hours post-transfection before fixation or live-cell imaging.

### Live-cell Imaging

Cells were grown to 80–90% confluency in 35-mm dishes with #1.5-thickness glass coverslip bottoms (MatTek, P35G-1.5-14-C). Imaging was performed on an Applied Precision OMX V4 microscope equipped with a 100×/1.4 NA oil immersion objective (Olympus), two Evolve EMCCDs (Photometrics), and a live-cell incubation chamber. The chamber maintained a humidified CO2 supply and temperature control at 37°C for both the incubator and the objective lens. Following drug treatment and HaloTag ligand addition as previously described, cells were acclimatized in the chamber for ∼1 hour. Imaging commenced 1–2 hours post-treatment, capturing 3D stacks (40–50 z-sections, 200 nm spacing) at 1-minute intervals for 1–2 hours. For each z-slice, we used simultaneous dual-camera acquisition. Nuclear speckles (EGFP-SON) and the HSPA1B BAC transgene (EGFP-LacR) were visualized using 488 nm excitation (2.0–10.0% transmittance, 5–10 ms exposure). HaloTag-labeled proteins were visualized using 642 nm excitation (5.0–10.0% transmittance, 8–10 ms exposure), adjusted to 10.0–31.0% transmittance and 8–15 ms exposure for HaloTag-TetR. mRFP-actin expression was assessed via a single pre-experiment timepoint using 568 nm excitation (2.0–10.0% transmittance, 8 ms exposure).

### Live-Cell Image Processing and Enhancement

For all frames in live-cell movies, 3D z-stacks were deconvolved using the Enhanced version of the iterative, nonlinear deconvolution algorithm provided by Softworx (Applied Precision, GE Healthcare). 2D projections were calculated using a maximum intensity projection algorithm. For live-cell movie analysis, individual nuclei were identified and the maximum intensity 2D projected area containing each nucleus was cropped and labeled with a hierarchical identifier based on the data collection multi-field position “ visit” number and nucleus number within that visit (e.g., v10n2, indicating visit 10, nucleus 2). A 3×3 pixel median filter was applied and followed by a Gaussian filter with a sigma radius of ∼0.8–1.6 pixels, adjusted based on image quality to minimize Poisson and camera noise.

To correct for nuclear rotation and translational movement, the StackReg plugin (version 2.0, Philippe Thévenaz, Biomedical Imaging Group, EPFL) for ImageJ/Fiji was applied using the “Rigid Body” transformation option. For C7MCP movies in which the *HSPA1B* BAC exhibited considerably higher fluorescent intensity from nuclear speckles, a gamma correction function was applied to decrease the overall intensity dynamic range; this rescaling prevented a disproportionate influence of the BAC spot-signal on the cross-correlation image alignment between adjacent time frames. For live-cell movies containing two fluorescent channels, image registration between adjacent time frames was performed using the HyperStackReg plugin (version 5.7, Ved Sharma, Albert Einstein College of Medicine, ImageJ/Fiji) with the “Rigid Body” transformation option. To ensure robust and unbiased registration, the transformation matrix was calculated using the reference channel containing the speckle marker (GFP-SON or HaloTag-DDX39B-Cy5) rather than the BAC channel. This transformation matrix was then applied to both channels simultaneously, ensuring accurate spatial alignment throughout the time series.

For nuclei exhibiting diffuse SON background signal and/or low speckle-to-background intensity ratios, a rolling ball background subtraction (rolling ball radius of approximately 20–50 pixels) was applied prior to Gaussian blur filtering to enhance speckle-to-background contrast. Finally, to correct for signal loss due to photobleaching over the course of the time-lapse movie, the BleachCorrect plugin (version 2.1.0, Kota Miura, ImageJ/Fiji^49^) was applied using either exponential fit or simple ratio correction methods, as appropriate for each image sequence.

### BAC position tracking and distance to speckle calculation

For live-cell imaging experiments, the position of the BAC locus and the nearest NS edge were manually tracked using the Manual Tracking plugin (version 2.1.1, Fabrice Cordelieres, Institut Curie) in ImageJ/Fiji to obtain x,y pixel coordinates at each time point. In cases where the BAC locus interacted with multiple NS, a secondary NS edge was also tracked. The distance between the BAC locus and the nearest NS edge was calculated at each frame using the Euclidean distance formula: 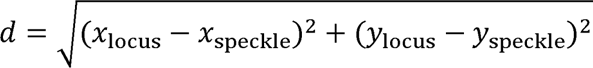, where *x*_locus_ and *y*_locus_ are the pixel coordinates of the gene locus, and *x*_speckle_ and *y*_speckle_ are the pixel coordinates of the NS edge. The calculated distance in pixels was then converted to micrometers using the calibration factor (0.08 µm/pixel). Distance values were plotted as a time series over the acquisition period (1 frame/minute). The “nearest” NS edge was defined as the NS showing the smallest distance to the gene locus when multiple NS were present.

The BAC locus mobility was assessed using the x,y coordinates converted to physical distances (µm) to calculate frame-to-frame displacement (µm) and velocity (µm/min). Step-size (the distance moved by the BAC locus between consecutive frames) was determined at each time point and averaged per locus to provide a measure of BAC mobility. Total BAC displacement over the entire movie was calculated and normalized to displacement per hour to account for cases in which there were variable movie lengths.

An oscillation was defined as the movement of a NS-associated gene locus away from a NS followed by its return and reattachment to either the same NS or a different NS. The start of an oscillation was identified as the frame in which the BAC locus was attached to the NS, immediately followed by detachment in the subsequent frame, evidenced by BAC displacement away from the speckle edge (BAC-NS separation >0.3 µm). The end of an oscillation was defined as the frame in which the BAC locus reattached to either the original NS or a different NS, evidenced by locus movement toward and reattachment to the NS (BAC-NS distance ≤0.3 µm).

Oscillations were quantified by visual inspection based on the 0.3 µm detachment threshold. Oscillations involving switches between nearby NS were included in the oscillation count, even in those cases in which the BAC locus never moved further than 0.3 µm away from the nearest NS; these cases were not captured in distance time-series analysis as the distance to the nearest NS would remain below 0.3 µm. In some cases, to visualize long-range movements, only oscillations with maximum BAC-NS separation ≥0.4 µm were displayed. In cases of low BAC mobility where the observable BAC trajectory displacement was limited, oscillation start and end frames were identified using time-series distance measurements to distinguish detached phases (BAC-NS distance >0.3 µm) from attached phases (BAC-NS distance ≤0.3 µm).

The oscillation frequency per nucleus was calculated as the total number of oscillations counted in each movie, normalized to oscillations per hour to account for variable movie lengths. For each identified oscillation, the start and end frames were used to calculate the following parameters: (1) **Oscillation duration** was calculated as the time elapsed (minutes) between the start and end frames, based on the acquisition rate of 1 frame/minute; (2) **Maximum distance from nuclear speckle** (d_max_, µm) was defined as the largest BAC-NS separation achieved during the oscillation, measured as the distance between the gene locus and the nearest NS edge; (3) **Total BAC displacement** (Δr, µm) was calculated as the sum of all frame-to-frame step distances accumulated throughout the oscillation, providing a quantitative measure of the total path length traveled by the *HSPA1B* BAC locus during each oscillation event.

### Confidence Interval Calculation for Stacked Bar Charts

Distance measurements from the HSPA1 locus or BAC transgene to nuclear speckles were binned into four non-overlapping distance categories (<0.3 μm, 0.3–0.45 μm, 0.45–0.6 μm, and ≥0.6 μm), with similar binning applied to gene-lamin distances (<0.7 μm, 0.7–1.5 μm, 1.5–2.5 μm, and ≥2.5 μm). Measurements were obtained across multiple experimental conditions including untreated controls and transcriptional inhibition treatments.

For each distance category, the percentage of observations and 95% confidence intervals (CI) were calculated using the Wilson Score interval method, which provides valid bounds [0%, 100%] for proportional data without boundary violations. The 95% CI lower and upper bounds were calculated using:

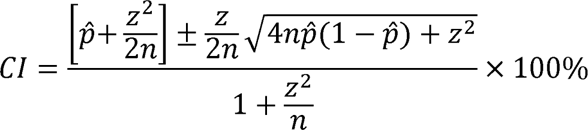

where *p*□ = observed proportion, *n* = sample size, and *z* = 1.96.

Error bars in stacked percentage bar charts were derived as CI_Upper - Percentage (upper) and Percentage - CI_Lower (lower). Overlapping error bars reflect sampling uncertainty in proportion estimates. Complete calculations and formula specifications are provided in the source data file.

### Temporal Color-Coded Trajectory Projection

To visualize gene locus movement paths and distinguish directional dynamics, temporal color-coded projections were applied to 2D time-lapse sequences. This technique assigns distinct colors from a lookup table (LUT) to sequential time frames, with early time points displayed in one end of the color spectrum and later frames progressively transitioning toward the opposite end. Overlapping pixels from multiple time frames combine to produce white coloration, highlighting stationary elements.

Three distinct LUT schemes were employed to visualize different movement characteristics: (1) The **Spectrum LUT** was used to display unidirectional movements and static trajectories, with stationary nuclear speckles or gene loci appearing in white and moving gene loci exhibiting color gradation; (2) The **Blue-Magenta-Red (BMR) LUT** was used to visualize gene locus movements away from nuclear speckles; (3) The **Cyan-Green-Yellow (CGY) LUT** was used to visualize gene locus movements toward nuclear speckles (both LUTs from Christophe Leterrier, ChrisLUTs collection, https://github.com/cleterrier/ChrisLUTs).

### Spatial Grid Generation for Trajectory Visualization

To further illustrate gene locus movement patterns, spatial trajectory grids were generated from individual movie frames or temporal color-coded projections. A spatial grid was constructed by overlaying circles at the gene locus position at multiple sequential time points, with each circle representing the locus location at a specific frame. Directional arrows were subsequently drawn between consecutive circles to indicate the vector of movement from one time point to the next, creating a simplified trajectory map. Grids were generated with appropriate dimensions corresponding to the nuclear region of interest, and overlays were created using Adobe Illustrator to ensure precise spatial registration and clear visual representation of locus movement directionality.

### Trajectory Measurement

Gene locus trajectories were visualized and measured using the line tool in Fiji/ImageJ. Individual trajectories consisted of unidirectional movements spanning 2–3 steps (3–4 positional points). For oscillatory movements involving detachment from and reattachment to nuclear speckles, separate linearity indices were calculated for the outward (away from speckle) and return (back to speckle) trajectories. For speckle-to-speckle transfers, a single linearity index was calculated for the unidirectional movement between the two speckles. Distance measurements between points and angles relative to the XY plane of the 2D image were recorded for each trajectory.

### Linearity Index Calculation

To quantify trajectory linearity, a linearity index was calculated as the ratio of total path length to net displacement straight-line distance from the start to the end position of the unidirectional movement. Where total path length is the sum of all frame-to-frame step distances, and net displacement is the straight-line distance from the start to the end position of the unidirectional movement. A linearity index of 1 indicates perfectly straight, unidirectional movement along the direct path. Values greater than 1 indicate increasingly curved or tortuous trajectories, with larger values reflecting greater deviations from the direct path.

### Angular Directionality Analysis and Linear/Curved Classification

For each trajectory segment, a reference vector was defined as the straight line connecting the start position to the end position of the movement. Individual step vectors were calculated for each frame-to-frame displacement, and the absolute angular offset (deviation) of each step vector relative to the reference vector was measured in degrees. The average angular offset across all steps within a trajectory was calculated. For classification purposes, trajectories were assigned as linear (average angular offset ≤25°) or curved (average angular offset >25°).

### Statistical Assessment of Trajectory Directionality

To test whether observed gene locus trajectories exhibit directional movement rather than random walk behavior, a Rayleigh test for uniformity was applied to the distribution of absolute angular offsets from the reference vector. Individual step vectors from all trajectories (n = 111 step vectors from 51 total trajectories) were pooled, and a Rayleigh test was applied using the circular package in R. The Rayleigh test evaluates the concentration of angles around the mean direction, calculating a mean resultant length R (range: 0–1, where values closer to 1 indicate stronger directional clustering). A p-value < 0.05 was considered statistically significant evidence that step vectors were non-randomly oriented, indicating directed rather than random movement.

### Step-to-Step Angular Deviations

Step-to-step angular deviations were calculated as the absolute difference in angle between consecutive step vectors within each trajectory. One-sample t-tests were performed separately for linear and curved trajectory groups to test whether observed deviations were significantly different from 0° (the expectation for perfectly linear movement). These tests were conducted in GraphPad Prism.

### Quantitative Assessment of Protein Levels via Fluorescence Microscopy for siRNA Knockdown

Quantitative measurement of protein localization was performed using ImageJ/FIJI software. The freehand area selection tool was used to carefully outline the nucleus, with borders defined by DAPI fluorescence in fixed cells or by GFP-SON speckle signal to identify the nucleoplasmic region in live cells. Mean gray intensity values were measured within the nuclear region of interest using the Cy5 channel to quantify signal from HaloTag-labeled proteins. To correct for non-specific background signal, a measurement region of equivalent size and shape was selected in the same image field outside the nucleus, and mean gray intensity was determined using identical acquisition settings. The relative protein enrichment in the nucleus compared to background was calculated using:

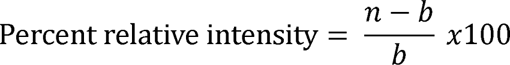

where *n* represents the mean gray value measured within the nucleus and *b* represents the mean gray value from the background region. This formula normalizes the nuclear signal to background intensity, yielding a percentage that reflects the degree of protein enrichment or depletion. Following successful knockdown of the target protein via siRNA, this value approaches zero or becomes negative, indicating loss of nuclear protein accumulation relative to control conditions.

## Supplementary Figure Legends

**Supplementary Figure 1. Inhibitors to RNA polymerase 2 and HSF1 decrease the association of the endogenous *HSPA1* locus in human HCT116 cells and an integrated human *HSPA1B* BAC transgene in CHO cells.**

(**a**) Violin plots showing distance distribution in fixed cells of the TetO-tagged endogenous *HSPA1* locus from the edge of nuclear speckles (NS) in human HCT116 cells (clone “96-8 B2”) under indicated treatment conditions at 37°C and after 30 min heat shock (HS). 37°C: n= 270, 96, 199, 249, 165, 174; HS: n= 223, 118, 179, 115, 347, 126 (No Treatment, DMSO, DRB, Triptolide,α-amanitin, KRIBB11, respectively). (**b**) Same as (**a**) but showing LacO-tagged *HSPA1B* BAC transgene distances in CHO K1 cells (clone “C7MCP”). 37°C: n= 421, 181, 283, 480, 293, 411; HS: n= 425, 430, 266, 195, 298, 281(No Treatment, DMSO, DRB, Triptolide,α-amanitin, KRIBB11, respectively).(**c**) Live-cell distance distributions of TetO-tagged HSPA1 endogenous locus from NS closely reproduce fixed-cell measurements at 37°C in untreated cells and after treatment with TPL or KRIBB11, indicating minimal phototoxicity-induced changes in HSPA1–NS association. Points represent individual measurements; red error bars show mean ±SD. Fixed cells: n = 270, 209, 174; live-cell measurements: n = 3,595, 3,014, 6,518 (No treatment, TPL, KRIBB11, respectively). Live-cell measurements were calculated using custom MATLAB code (see Methods). **For (a), (b) and (c) :** Kruskal-Wallis test with Dunn’s multiple comparisons test; ns, P > 0.05; *P ≤ 0.05; **P ≤ 0.01; ***P ≤ 0.001; ****P ≤ 0.0001. (**d**) Number of oscillations per hour to and from NS where gene–NS separation reaches at least 0.4 μm. Top: endogenous HSPA1 locus in HCT116 cells. Bottom: HSPA1B BAC transgene in CHO C7MCP. Both measured at 37°C under control conditions (No Treat.) or after treatment with DRB, TPL, or α-amanitin. Bar height shows mean ±SD. HCT116: n = 7, 5, 6, 5; CHO: n = 9, 8, 10, 10 nuclei (No Treat., DRB, TPL, α-amanitin, respectively). (**e**) Oscillation characteristics: maximum gene–NS separation distance and movement type. Left: Percentage of oscillations within indicated maximum distance ranges from NS. Right: Percentages of oscillation movements between NS (green) versus to/from a single NS (blue) after no treatment or treatment with DRB, TPL, or α-amanitin. Top: endogenous HSPA1 locus in HCT116 cells. Bottom: HSPA1B BAC transgene in CHO K1 cells (clone C7MCP). (**f**) Representative examples of oscillation dynamics in untreated cells. Top: Endogenous HSPA1 locus (red) maintains nearly stable NS association (green) with minimal gene–NS separation, as shown by distance measurements (middle) and individual time-frame images (right) at color-coded timepoints. Bottom: Same as top but for HSPA1B BAC transgene (bright white) relative to NS (grey).

**Supplementary Figure 2. Comparisons of total displacement versus maximum distance from nuclear speckles per oscillation consistent with mostly curvilinear *HSPA1* gene and transgene movements**

(**a-b**) maximum BAC–nuclear speckle separation distance (x-axis) versus total BAC displacement (y-axis; Δr) for endogenous *HSPA1* (HCT116 cells; **a**) and *HSPA1B* BAC (CHO C7MCP cells; **b**) under transcriptional inhibitor treatments. Bubble color indicates oscillation duration (minutes). n = 54, 55, 123, 123 oscillations from 7, 5, 6, 5 nuclei (endogenous *HSPA1*) and n = 45, 152, 234, 265 oscillations from 7, 8, 10, 10 nuclei (*HSPA1B* BAC) for No Treatment, DRB, Triptolide, and α-amanitin, respectively.

**Supplementary Figure 3. Deletion of *HSPA1* genes and flanking DNA sequence from *HSPA1* BAC reduces both nuclear speckle association and curvilinear oscillations**

(**a**) Schematic of full-length *HSPA1* BAC transgene (WT; top) containing the human HSPA1 locus, versus the *HSP*Δ (“D3”) deletion BAC lacking all three *HSPA1* genes (middle), and the *HSP*+EnhRg Δ (“HSeD”) BAC (bottom) including the D3 deletion plus additional 3’ flanking DNA corresponding to a region enrighed for H3K27ac at the endogenous locus in fibroblasts. (**b-d**) Distance measurements in fixed cells without treatment (No Treat.) or after triptolide (TPL) showing BAC–nuclear speckle (NS) distances (**b,d**, left) and BAC–nuclear lamina distances (**c,d**, right) for three cell lines containing WT, *HSP*Δ, or *HSP*+EnhΔ BACs. Data are shown as mean ± SD, with each point representing one BAC transgene. **(b)** n = 113, 60 (WT); 228, 133 (*HSP*Δ); and 121, 174 (*HSP*+EnhΔ) loci for No Treat. and TPL, respectively. **(c)** n = 64, 29 (WT); 92, 51 (*HSP*Δ); and 121, 174 (*HSP*+EnhΔ) loci for No Treat. and TPL, respectively. **(d)** Same datasets as in **(b & c)** plotted as the percentage of loci within specified BAC–NS or BAC–lamina distance ranges. Error bars represent 95% confidence intervals calculated using the Wilson Score method (see Methods for details). Overlapping error bars reflect sampling uncertainty in proportion estimates. (**e-g**) Representative live-cell examples after TPL treatment comparing curvilear movements of the WT BAC transgene (HaloTag-Tet repressor, red) (**e**) away from NS (EGFP-SON, green) at 15-17 min and back at 18-20 min, with the more random and relatively stationary behaviour of the *HSP*Δ (**f**) and *HSP*+Enh Δ (**g**) BACs. Color-coded temporal projections for the indicated time windows show that the WT BAC trajectories returning to the NS largely retrace paths taken away from the NS (**e**, right), whereas deletion BAC trajectories appear more stationary and dispersed over longer intervals (**f,g**, right). (**h**) Oscillation frequency (events/hour, any BAC–NS distance range) in the *HSPA1B* BAC cell line compared with WT and deletion BAC lines. Data are mean ± SD; n = 10 nuclei (*HSPA1B* BAC) and n = 5 nuclei per *HSPA1* WT or deletion BAC. **(i)** Comparisonof BAC transgene dynamics from live-cell movies for three nuclei (Nuc #1–3; corresponding to **e–g**) in *HSPA1* BAC WT and deletion cell lines acquired at 1 frame/min. Left, BAC–NS distance in each frame (mean ± SD). Right, BAC step size between adjacent frames (bar height = mean ± SD); each point represents an individual frame-to-frame measurement. Frames: n = 121, 66 and 91 for WT, *HSP*Δ and *HSP*+EnhΔ, respectively. **(b, c, i)** Kruskal–Wallis test with Dunn’s multiple comparisons test was used. P-value notation: ns (P > 0.05), *P ≤ 0.05, **P ≤ 0.01, ***P ≤ 0.001, ****P ≤ 0.0001.

**Supplementary Figure 4. KRIBB11 inhibition of HSF1 disrupts directional movement and induces random transgene trajectories**

**(a**) Scatter plot of individual BAC step sizes (µm/min) per treatment. Each point represents one step; data shown as mean ± SD. Steps: n = 936, 960, 1170, 1179 and 239 from 9, 8, 10, 10 and 3 nuclei for No treatment, DRB, triptolide (TPL), α-amanitin and KRIBB11, respectively. (**b**) Individual oscillation statistics for *HSPA1B* BAC after KRIBB11 treatment. Total BAC displacement (sum of frame-to-frame steps, y-axis) plotted against maximum distance from speckle edge (µm, x-axis). Color indicates oscillation duration (minutes; color bar) at 1 frame/min acquisition (n= 22 oscillations from 3 nuclei). (**c**) Oscillation frequency (events/hour) per treatment. Each point represents one nucleus; error bars show mean ± SD. Top, all oscillation events. Bottom, subset of oscillations in which BAC-NS separation eached ≥0.4 µm. n=9, 8, 10, 10, 3 nuclei for No treatmntm DRB, TPL, α-amanitin and KRIBB11, respectively. (**d**) Time series of BAC-speckle distance for representative nuclei after TPL (left) or KRIBB11 (right) at 1 frame/min. Orange shading from 0–0.4 µm denotes the speckle-associated zone used as a reference for oscillation start and end. Right, spatial grids of BAC trajectories corresponding to labeled time points. TPL treatment shows transgene path retracing and return to start position, whereas KRIBB11 produces more random, non-retracing motion. Temporal color-coded projection of frames within oscillation “A” (KRIBB11, right) reveals BAC transgene remaining relatively stationary for multiple frames during movement to and from the speckle. (**e**) Quantification of oscillation frequency (events/hour) and oscillation duration (minutes) after TPL versus KRIBB11 for 3 nuclei per condition. Measurements are relatively consistent among TPL-treated nuclei but vary widely between KRIBB11-treated nuclei. **(f)** Scatter plots of HSF1 knockdown efficiency, measured as percentage mean nuclear HaloTag–HSF1 intensity over background for fixed cells (left) and live-cell imaged nuclei (right). Each point represents one nucleus under No treatment (grey triangles) or after TPL (blue circles), transfected with NC-1 control siRNA or HSF1 siRNA. Data are mean ± SD. Fixed cells: NC-1 n = 44 (No treatment) and 136 (TPL); HSF1 siRNA n = 104 (No treatment) and 173 (TPL). Live cells: NC-1 n = 95 (No treatment); HSF1 siRNA n = 111 (No treatment) and 49 (TPL). **(g)** Scatter plot of BAC–NS distance in fixed cells under No treatment (grey triangles) or after TPL (blue circles). n = 35 and 104 nuclei (No treatment), and 122 and 173 nuclei (TPL) for NC-1 and HSF1 siRNA, respectively; data are mean ± SD. **(a, f, g)** Kruskal–Wallis test with Dunn’s multiple comparisons test was used. P-value notation: ns (P > 0.05), *P ≤ 0.05, **P ≤ 0.01, ***P ≤ 0.001, ****P ≤ 0.0001.

**Supplementary Figure 5. HSPA1B BAC locus oscillations and long-range motility persist following actin filament disruption**

**(a)** The experimental sequence (transcriptional inhibitor → actin/myosin inhibitor → fixation) tests whether actin filament disruption prevents BAC transgene movement triggered by transcriptional activation. Representative immunofluorescence images from untreated and triptolide-treated CHO cells showing actin filament architecture following perturbation with actin/myosin inhibitors. Top row, untreated control cells (No Treatment). Bottom row, cells pre-treated with Triptolide (TPL) to trigger BAC transgene movement, followed by addition of latrunculin A (LatA) or cytochalasin D (CytoD) before fixation. Images display *HSPA1B* BAC locus (EGFP-LacI fusion protein, green), nuclear speckles (NS) (EGFP-SON, green), DNA (DAPI, blue), and actin filaments (phalloidin staining, Cy5, red). (**b**) Scatter plot of *HSPA1B* BAC distance to NS edge from fixed cells under control conditions (gray triangles) or after 2 hours TPL treatment (blue circles), without additional inhibitor, with myosin VI inhibitor (triiodophenol, TIP), or with actin inhibitors (CytoD or LatA). Sample sizes: n = 55, 76, 30, 89, 38, 71, 44, 61 for No Treatment/Control, TPL/Control, No Treatment/TIP, TPL/TIP, No Treatment/CytoD, TPL/Cyto D, No Treatment/LatA, and TPL/LatA, respectively. Each point represents one cell; red line indicates mean ± SD. (**c**) Scatter plot of *HSPA1B* BAC distance to NS edge in fixed cells following 2 hours TPL treatment. Cells were either untransfected (control) or transfected with mRFP-NLS-actin constructs: wildtype (WT), S14C mutant, or G13R mutant. Sample sizes: n = 480 (no plasmid), 56 (WT actin), 61 (S14C actin), 99 (G13R actin). Each point represents one cell; line indicates mean ± SD. (**d**) Live-cell imaging of cells transfected with mRFP-actin plasmids expressing either G13R or S14C mutant actin. Top, snapshot before live-cell imaging showing mRFP signal (red) demonstrating G13R actin plasmid (left) or S14C actin plasmid (right) expression, along with *HSPA1B* BAC locus (EGFP-LacI, green) and nuclear speckles (EGFP-SON, green). Middle, grayscale maximum-intensity projection of *HSPA1B* BAC and speckle channels from live-cell time series. Bottom, temporal color-coded projection of multiple frames showing long-range BAC transgene movement toward or away from nuclear speckles, with corresponding spatial grid indicating BAC position and movement as shown in projection. **e,** Grayscale image of CHO C7MCP cell expressing S14C actin plasmid showing long-range movement of *HSPA1B* BAC transgene across nucleus over approximately 20 minutes. Right, temporal color-coded projection of frames using spectrum lookup table (LUT); white arrow indicates BAC movement direction over the projection. (**f**) Live-cell movie frames demonstrating *HSPA1B* BAC transgene movement dynamics persist after actin perturbation. Left, representative frames showing BAC transgene movement away from NS (red arrow) and return to NS (green arrow) after DRB treatment. Middle, time-series plot of BAC distance to speckle from live-cell movies of the same cell following DRB treatment and subsequent addition of latrunculin A (LatA). Right, representative frames showing oscillatory BAC movement persists despite actin filament disruption by LatA treatment. Frame time points correspond to red or green highlights in the time-series plot. Acquisition rate: 1 frame per 30 seconds. Statistical significance determined by Kruskal-Wallis test with Dunn’s multiple comparisons. Significance thresholds: ns (p > 0.05), *p ≤ 0.05, **p ≤ 0.01, ***p ≤ 0.001, ****p ≤ 0.0001.

**Supplementary Figure 6.**

Live-cell imaging of *HSPA1B* BAC transgene dynamics in three nuclei per condition following TPL treatment with additional TPL (control), myosin IV TIP inhibitor, or cytochalasin D inhibitor. Time-lapse acquisition at 1 frame/minute: 80 minutes for TPL+TPL and TPL+TIP conditions, 120 minutes for TPL+CytoD condition**. a, Left,** Scatter plot of BAC distance to nearest speckle edge from time-series measurements (n = 243, 243, 363 timepoints for TPL+TPL, TPL+TIP, and TPL+CytoD, respectively; each point represents one measurement). **a, Right**, Scatter plot of BAC step size (µm/min) from same time-series analysis (n = 240, 240, 360 steps, respectively; each point represents one measured step). **b, Left,** Oscillation count per nucleus normalized to oscillations/hour for three nuclei per treatment (each point represents one nucleus). **b, Right**, BAC transgene dynamics measurements pooled from all three nuclei per treatment showing maximum distance BAC reaches from speckle per oscillation and total BAC displacement per oscillation (sum of steps; each point represents one oscillation; n = 53, 55, 83). All scatter plots display mean (red line) ± SD. **c,** Measurements from fixed C7MCP *HaloTag-RAD21* knockin cells transfected with siRNA targeting *RAD21* (siRNA-1) or negative control siRNA (NC-1), either untreated (gray triangles) or treated with TPL (blue circles). Each point represents one cell. Left, *RAD21* protein level measured by fluorescence intensity (percent over background) to assess knockdown efficiency in NC-1 and *RAD21* siRNA-1 cells (n = 24, 28, 24 for untreated NC-1, untreated *RAD21* siRNA-1, and TPL-treated *RAD21* siRNA-1, respectively). Right, Distance of *HSPA1B* BAC transgene to nearest nuclear speckle (n = 50, 53, 37, 39 for siRNA NC-1 untreated, NC-1 + TPL, *RAD21* siRNA-1 untreated, and *RAD21* siRNA-1 + TPL, respectively). All scatter plots display mean (red line) ± SD. **d,** *HSPA1B* BAC transgene oscillation dynamics in TPL-treated cells. Gray triangles represent previously measured TPL-treated control cells; green squares represent cells following *RAD21*siRNA-1 transfection. Left, Oscillation count normalized to oscillations/hour for 10 nuclei in control and 3 nuclei after *RAD21* siRNA-1 transfection. Right, Individual oscillation dynamics showing maximum distance BAC reaches from speckle and total BAC displacement per oscillation, pooled from 10 (control) or 3 (*RAD21* siRNA-1) nuclei (n = 234 and 57 oscillations, respectively; each point represents one oscillation). All scatter plots display mean (red line) ± SD. **e,** Frames from live-cell movie of *RAD21* siRNA-transfected cells with TPL treatment. Left (Nuc #1), Representative *HSPA1B* BAC transgene exhibiting near-linear long-range motion between nuclear speckles, displayed using spectrum lookup table (LUT) to visualize movement trajectory from one nuclear speckle to another and directional motion toward nuclear speckles. Corresponding spatial grid of BAC position and movement trajectory is shown to the right. Right (Nuc #2), Representative oscillatory *HSPA1B* BAC movement showing frames of BAC motion toward nuclear speckles with corresponding spatial grid, as well as additional spatial grids below displaying overlapping oscillatory BAC movement trajectories. Statistical significance determined by Kruskal-Wallis test with Dunn’s multiple comparisons. Significance thresholds: ns (p > 0.05), *p ≤ 0.05, **p ≤ 0.01, ***p ≤ 0.001, ****p ≤ 0.0001.

**Supplementary Figure 7. D*D*X39B knockdown leads to chromatin reorganization, SON-marked speckle dispersion, and apoptotic cell death.**

In all panels, green (FITC) shows nuclear speckles (EGFP-SON) and the *HSPA1B* BAC transgene (EGFP-LacI), and red (Cy5) shows HaloTagged *DDX39B*; in panel c, blue (DAPI) marks DNA. **a**, Frames from live-cell imaging of a CHO C7MCP cell after TPL treatment at 37 °C, showing *HSPA1B* BAC and SON-marked speckles with HaloTagged *DDX39B*. *DDX39B* forms bridges connecting non-speckle-associated BACs to speckles, and small *DDX39B* droplets contact immobile, non-speckle-associated BACs during periods of BAC immobility. **b**, Frames from live-cell imaging of a CHO C7MCP cell transfected with *DDX39B* siRNA-2 and after TPL treatment at 37°C. **Left**, Six representative frames from a 61-frame movie (60 min total, 1 frame/min acquisition) showing dispersal and disappearance of SON-marked speckles followed by speckle re-nucleation over time, while the *HSPA1B* BAC transgene remains relatively immobile. BAC transgene locus is outlined with a dashed orange square to distinguish it from speckles. **Right**, Temporal projection of all 61 frames using spectrum lookup table (LUT). The BAC region is outlined with a dashed orange line; overlapping BAC positions create white signal, while varied coloration throughout the nucleus represents speckle dispersal and re-nucleation dynamics. **c**, Fixed CHO C7MCP cells 38 hours post-transfection with siRNA targeting *DDX39B* or negative control. **Top**, NC-1 negative control and *DDX39B* siRNA-2. **Bottom**, *DDX39B* siRNA-3. Untreated cells at 37 °C. Maximum intensity projection of 11, 12, or 14 z-sections with 200 nm spacing. *DDX39B* knockdown leads to apparent chromatin reorganization, SON-marked nuclear speckle dispersion, and apoptotic nuclear morphology.

## Supplementary Information

**Movie 1: Transcriptional inhibition induces oscillatory HSPA1 endogenous gene and *HSPA1B* transgene movements to and from nuclear speckles and between nuclear speckles**

Live-cell movies acquired at 37 °C with 1 frame/min, comparing untreated cells (left) to cells after triptolide treatment (right). Top: two nuclei from HCT116 96-8 B2 cells with endogenous SON (EGFP, green) and endogenous *HSPA1* (HaloTag-Cy5, red), corresponding to Fig. 1c and SFig. 1f. Bottom: two nuclei from CHO C7MCP cells showing nuclear speckles (EGFP-SON BAC, gray) and the *HSPA1B* BAC transgene (bright gray), corresponding to Fig. 1g and SFig. 1f. Yellow boxes highlight tagged loci/transgene and interacting speckles. White upper-left timestamps indicate movie frames (hr:min); green upper-right timestamps mark time after triptolide (hr:min). Scale bar, 2 µm.

**Movie 2: Closer view of oscillatory *HSPA1B* transgene movements to and from nuclear speckles induced after triptolide or alpha-amanitin transcriptional inhibition**

Live-cell movie of CHO C7MCP cells at 37 °C with 1 frame/minute acquisition, comparing triptolide treatment (top three nuclei) with α-amanitin treatment (bottom three nuclei). Nuclear speckles (EGFP-SON BAC, gray) and the *HSPA1B* BAC transgene (gray) are visualized. Red box highlights the BAC transgene every 10 minutes. Top left timestamp indicates movie frame time (minutes). Bottom right timestamp indicates treatment time (hr:min): green for triptolide, orange for α-amanitin. For triptolide-treated cells: left, center, and right movies correspond to Fig. 2a, 2b, and 2d, respectively. For α-amanitin-treated cells: left nucleus corresponds to Fig. 2b and 2e; center nucleus to Fig. 2d.

**Movie 3: KRIBB11 treatment reduces BAC transgene movements and oscillations and results in more random gene trajectories and increased nuclear speckle detachment**

Live-cell movie of three nuclei from CHO C7MCP cells imaged at 37 °C with 1 frame/minute acquisition following KRIBB11 treatment. Left panel corresponds to nucleus shown in Fig. 3a-c. Nuclear speckles (EGFP-SON BAC) and *HSPA1B* BAC transgene are both shown in gray. Every 10 minutes, a red box highlights the BAC transgene. Upper-right label shows time (minutes), matching frames displayed in Fig. 3. Lower-left orange time stamp indicates KRIBB11 treatment time (hr:min). Scale bar = 2 µm.

**Movie 4: *HSF1* knockdown results in reduced *HSPA1B* BAC movement and more random gene trajectories**

Live-cell movies of five nuclei contrast behavior of BAC transgene in cells with significant siRNA knockdown of HSF1 (white labels, Nuclei (Nuc) #2,4,5) versus in cells that retain high HSF1 levels (red labels, Nuclei (Nuc) #1,3). Imaging was performed after triptolide treatment with imaging at 37 °C using 1 minute intervals. Labeling of nuclei is the same as in Fig. 3d-h. Nuclear speckles (EGFP-SON BAC) and *HSPA1B* BAC transgene are both shown in gray-scale. Red box highlights BAC transgene location every 5 minutes. Time stamps (upper left panel) show data acquisition time (minutes, white) and time after addition of triptolide (hours:min, green). Scale bar = 1 µm.

**Movie 5: *HSPA1B* BAC transgene long-range movements and oscillations to and from nuclear speckles continue after *RAD21* knockdown**

Live-cell movies of three nuclei from CHO C7MCP cells following *RAD21* siRNA transfection and triptolide treatment, imaged at 37 °C with 1 frame/minute acquisition. Nuclear speckles (EGFP-SON BAC, gray) and the *HSPA1B* BAC transgene (gray) are shown. Every 5 minutes, a red box highlights the *HSPA1B* BAC transgene to distinguish it from the nuclear speckles. Nuclei are labeled Nuc #1–3 corresponding to frames in SFig. 6e. Upper-right timestamps indicate movie time (minutes), matching frames displayed in SFig. 6. Lower-right green timestamp marks triptolide treatment time (hr:min). Scale bar = 2 µm.

**Movie 6: Spatial and temporal correlation between *HSPA1B* BAC transgene movements and DDX39B protrusions and retractions from nuclear speckles**

Two movies showing examples of the correlation between *HSPA1B* BAC transgene movements and DDX39B condensate dynamics in triptolide-treated cells showing HaloTag–DDX39B (left panels, Cy5, red), EGFP-SON-marked nuclear speckles plus the brighter HSPA1B BAC transgene in green (EGFP-Lac repressor) (middle panels) and the merged images (right panels). (a) Nucleus shown corresponds to Fig. 4a. (b) Nucleus shown corresponds to Fig. 4b. Time stamps: top left (white, mins) shows data acquisition time as in Fig. 4 and top right (green, hours:minutes) shows time since triptolide addition. Scale bar = 2 µm.

**Movie 7: DDX39B siRNA knockdown reduces HSPA1B BAC transgene motility and increases BAC distance to nuclear speckles**

Live-cell imaging at 37°C following triptolide treatment and DDX39B knockdown at time intervals of one minute shows two examples. (a) BAC-speckle association/dissociation driven by speckle movement rather than BAC motility-corresponds to Fig. 5e (Nuc#4) (19% DDX39B mean intensity over background, siRNA-3). (b) BAC remains stationary; nearby small nuclear speckle shows nucleation and dispersion. (73% DDX39B mean intensity over background, siRNA-2). Right panels show HSPA1B BAC (EGFP-lac repressor) and nuclear speckles (EGFP-SON) (green, top) and DDX39B-HaloTag (red, bottom). Left shows merged channels with yellow box highlighting BAC position at 10-minute intervals. Time stamps: white (lower right) shows time (min) of data acquisition, green (upper left) shows time since triptolide addition (hr:min). Scale bar = 2 µm. Note: selection of nuclei for DDX39B KD was done based on mean intensity over background of 10–80% for DDX39B siRNA-transfected cells, and 70–115% for NC-1 controls.

## Acknowledgements

This work was supported by the National Institute of General Medical Sciences grant R01GM058460 (A.S.B) and by the National Institutes of Health Common Fund 4D Nucleome Program grants UM1HG011593 and U01DK127422 (A.S.B). We acknowledge the DNA Services and High-Performance Computing in Biology (HPCBio) group at the Roy J. Carver Biotechnology Center, University of Illinois at Urbana-Champaign, with particular thanks to Kimberly Walden, for assistance with sequencing and bioinformatic analysis of the CHO C7MCP cell line BAC integration.

## Data Availability

Data deposition to the Illinois Data Repository is in progress.

## Author Contributions

G.H.G. and A.S.B. conceived and designed the study based on preliminary data and observations provided by J.K. G.H.G. performed and analyzed the majority of experiments, including cell line generation via CRISPR knockin, fixed cell sample preparation and analysis, live-cell imaging, BAC mobility analysis, oscillation dynamics analysis, linearity and directionality analysis, and spatial grid generation. J.K. performed analysis of HCT116 live-cell imaging experiments using MATLAB. G.H.G. and P.G. contributed to cell line generation via CRISPR knockin of HaloTag-DDX39B and -RAD21, and G.H.G., P.G., and S.M. contributed to the HaloTag-HSF1 cell line generation and assisted in associated experiments. G.H.G. and P.G. performed RAD21 RNAi live-cell imaging experiments, and P.G. performed the associated fixed-cell experiments and analysis. P.C. provided guidance and contributed to BAC recombineering. P.K. performed TSA-seq experiments for the CHO K1 cell line. N.V. shared unpublished results about DDX39B leading to our correlation of DDX39B and HSPA1 dynamics.

## References

1. van Steensel, B. & Belmont, A. S. Lamina-Associated Domains: Links with Chromosome Architecture, Heterochromatin, and Gene Repression. Cell 169, 780–791 (2017).

2. Belmont, A. S. Nuclear Compartments: An Incomplete Primer to Nuclear Compartments, Bodies, and Genome Organization Relative to Nuclear Architecture. Cold Spring Harb Perspect Biol 14, (2022).

3. Bickmore, W. A. The spatial organization of the human genome. Annu Rev Genomics Hum Genet 14, 67–84 (2013).

4. Kumar, Y., Sengupta, D. & Bickmore, W. A. Recent advances in the spatial organization of the mammalian genome. Journal of Biosciences 45, (2020).

5. Misteli, T. The Self-Organizing Genome: Principles of Genome Architecture and Function. Cell 183, 28–45 (2020).

6. Takizawa, T., Meaburn, K. J. & Misteli, T. The meaning of gene positioning. Cell 135, 9–13 (2008).

7. Ferrai, C., de Castro, I. J., Lavitas, L., Chotalia, M. & Pombo, A. Gene positioning. Cold Spring Harb Perspect Biol 2, a000588 (2010).

8. Willemin, A., Szabó, D. & Pombo, A. Epigenetic regulatory layers in the 3D nucleus. Molecular Cell 84, 415–428 (2024).

9. Hall, L. L., Smith, K. P., Byron, M. & Lawrence, J. B. Molecular anatomy of a speckle. Anat Rec A Discov Mol Cell Evol Biol 288, 664–75 (2006).

10. Spector, D. L. & Lamond, A. I. Nuclear speckles. Cold Spring Harb Perspect Biol 3, (2011).

11. Galganski, L., Urbanek, M. O. & Krzyzosiak, W. J. Nuclear speckles: molecular organization, biological function and role in disease. Nucleic Acids Res 45, 10350–10368 (2017).

12. Chen, Y. et al. Mapping 3D genome organization relative to nuclear compartments using TSA-Seq as a cytological ruler. J Cell Biol 217, 4025–4048 (2018).

13. Gholamalamdari, O. et al. Major nuclear locales define nuclear genome organization and function beyond A and B compartments. eLife 13, RP99116 (2025).

14. Khanna, N., Hu, Y. & Belmont, A. S. HSP70 transgene directed motion to nuclear speckles facilitates heat shock activation. Curr Biol 24, 1138–44 (2014).

15. Kim, J., Venkata, N. C., Hernandez Gonzalez, G. A., Khanna, N. & Belmont, A. S. Gene expression amplification by nuclear speckle association. J Cell Biol 219, (2020).

16. Zhang, L. et al. TSA-seq reveals a largely conserved genome organization relative to nuclear speckles with small position changes tightly correlated with gene expression changes. Genome Res https://doi.org/10.1101/gr.266239.120 (2020) doi:10.1101/gr.266239.120.

17. Alexander, K. A. et al. p53 mediates target gene association with nuclear speckles for amplified RNA expression. Mol Cell 81, 1666–1681 e6 (2021).

18. Yu, R. et al. CTCF/RAD21 organize the ground state of chromatin-nuclear speckle association. Nat Struct Mol Biol 32, 1069–1080 (2025).

19. Venkata, N. C. et al. Highly active chromosome regions preferentially associate with two perispeckle networks that partition the interchromatin space. 2025.08.23.671945 Preprint at 10.1101/2025.08.23.671945 (2025).

20. Marshall, W. F. et al. Interphase chromosomes undergo constrained diffusional motion in living cells. Curr Biol 7, 930–9 (1997).

21. Chubb, J. R., Boyle, S., Perry, P. & Bickmore, W. A. Chromatin motion is constrained by association with nuclear compartments in human cells. Curr Biol 12, 439–45 (2002).

22. Chuang, C.-H. & Belmont, A. S. Moving chromatin within the interphase nucleus-controlled transitions? Seminars in Cell & Developmental Biology 18, 698–706 (2007).

23. Mazzocca, M., Narducci, D. N., Grosse-Holz, S., Matthias, J. & Hansen, A. S. Chromatin Dynamics are Highly Subdiffusive Across Seven Orders of Magnitude. 2025.05.10.653248 Preprint at 10.1101/2025.05.10.653248 (2025).

24. Hu, Y., Kireev, I., Plutz, M. J., Ashourian, N. & Belmont, A. S. Large-scale chromatin structure of inducible genes-transcription on a linear template. J. Cell Biol. 185, 87–100 (2009).

25. Hu, Y., Plutz, M. & Belmont, A. S. Hsp70 gene association with nuclear speckles is Hsp70 promoter specific. J Cell Biol 191, 711–9 (2010).

26. Hu, Y. Changes in large-scale chromatin structure and dynamics of BAC transgenes during transcriptional activation. (University of Illinois at Urbana-Champaign, 2010).

27. Tasan, I. et al. CRISPR/Cas9-mediated knock-in of an optimized TetO repeat for live cell imaging of endogenous loci. Nucleic Acids Res 46, e100 (2018).

28. Hu, Y. Changes in large-scale chromatin structure and dynamics of BAC transgenes during transcriptional activation. Cell and Developmental Biology vol. Ph.D (University of Illinois at Urbana-Champaign, 5.142010).

29. Nguyen, V. T. et al. In vivo degradation of RNA polymerase II largest subunit triggered by alpha-amanitin. Nucleic Acids Res 24, 2924–9 (1996).

30. Chuang, C. H. et al. Long-range directional movement of an interphase chromosome site. Curr Biol 16, 825–31 (2006).

31. Zhang, H. et al. Reversible phase separation of HSF1 is required for an acute transcriptional response during heat shock. Nat Cell Biol 24, 340–352 (2022).

32. Pincus, D. et al. Genetic and epigenetic determinants establish a continuum of Hsf1 occupancy and activity across the yeast genome. Mol Biol Cell 29, 3168–3182 (2018).

33. Vihervaara, A. et al. Transcriptional response to stress in the dynamic chromatin environment of cycling and mitotic cells. Proceedings of the National Academy of Sciences 110, E3388–E3397 (2013).

34. Yoon, Y. J. et al. KRIBB11 inhibits HSP70 synthesis through inhibition of heat shock factor 1 function by impairing the recruitment of positive transcription elongation factor b to the hsp70 promoter. J Biol Chem 286, 1737–1747 (2011).

35. Dundr, M. et al. Actin-dependent intranuclear repositioning of an active gene locus in vivo. J Cell Biol 179, 1095–103 (2007).

36. Wang, A. et al. Mechanism of Long-Range Chromosome Motion Triggered by Gene Activation. Dev Cell 52, 309–320 e5 (2020).

37. Hari-Gupta, Y. et al. Myosin VI regulates the spatial organisation of mammalian transcription initiation. Nat Commun 13, 1346 (2022).

38. Chen, Y. & Belmont, A. S. Genome organization around nuclear speckles. Curr Opin Genet Dev 55, 91–99 (2019).

39. Posern, G., Miralles, F., Guettler, S. & Treisman, R. Mutant actins that stabilise F-actin use distinct mechanisms to activate the SRF coactivator MAL. Embo J 23, 3973–83 (2004).

40. Fujiwara, I., Zweifel, M. E., Courtemanche, N. & Pollard, T. D. Latrunculin A Accelerates Actin Filament Depolymerization in Addition to Sequestering Actin Monomers. Current Biology 28, 3183–3192.e2 (2018).

41. Heissler, S. M. et al. Kinetic properties and small-molecule inhibition of human myosin-6. FEBS Lett 586, 3208–3214 (2012).

42. Balachandran, R. S. et al. The ubiquitin ligase CRL2ZYG11 targets cyclin B1 for degradation in a conserved pathway that facilitates mitotic slippage. J Cell Biol 215, 151–166 (2016).

43. Kim, J., Gonzalez, G. A. H., Venkata, N. C., Han, K. Y. & Belmont, A. S. Nonrandom interchromatin trafficking through dynamic multiphase speckle connections. 2025.05.23.655761 Preprint at 10.1101/2025.05.23.655761 (2025).

44. Strom, A. R. et al. Condensate interfacial forces reposition DNA loci and probe chromatin viscoelasticity. Cell 187, 5282–5297.e20 (2024).

45. Chaturvedi, P. et al. Acidic transcription factors position the genome at nuclear speckles through transcription dependent and independent mechanisms. 2025.10.01.679912 Preprint at 10.1101/2025.10.01.679912 (2025).

46. Yu, Y. et al. An efficient gene knock-in strategy using 5′-modified double-stranded DNA donors with short homology arms. Nat Chem Biol 16, 387–390 (2020).

47. Warming, S., Costantino, N., Court, D. L., Jenkins, N. A. & Copeland, N. G. Simple and highly efficient BAC recombineering using galK selection. Nucleic Acids Res 33, e36 (2005).

48. Kim, J., Han, K. Y., Khanna, N., Ha, T. & Belmont, A. S. Nuclear speckle fusion via long-range directional motion regulates speckle morphology after transcriptional inhibition. J Cell Sci 132, (2019).

49. Miura, K. Bleach correction ImageJ plugin for compensating the photobleaching of time-lapse sequences. Preprint at 10.12688/f1000research.27171.1 (2020).

