## Supplementary figures and images for "HSF1-dependent, long-range directional *HSPA1* gene motion to nuclear speckles is coupled to DDX39B condensate dynamics"

### Supplementary Figures

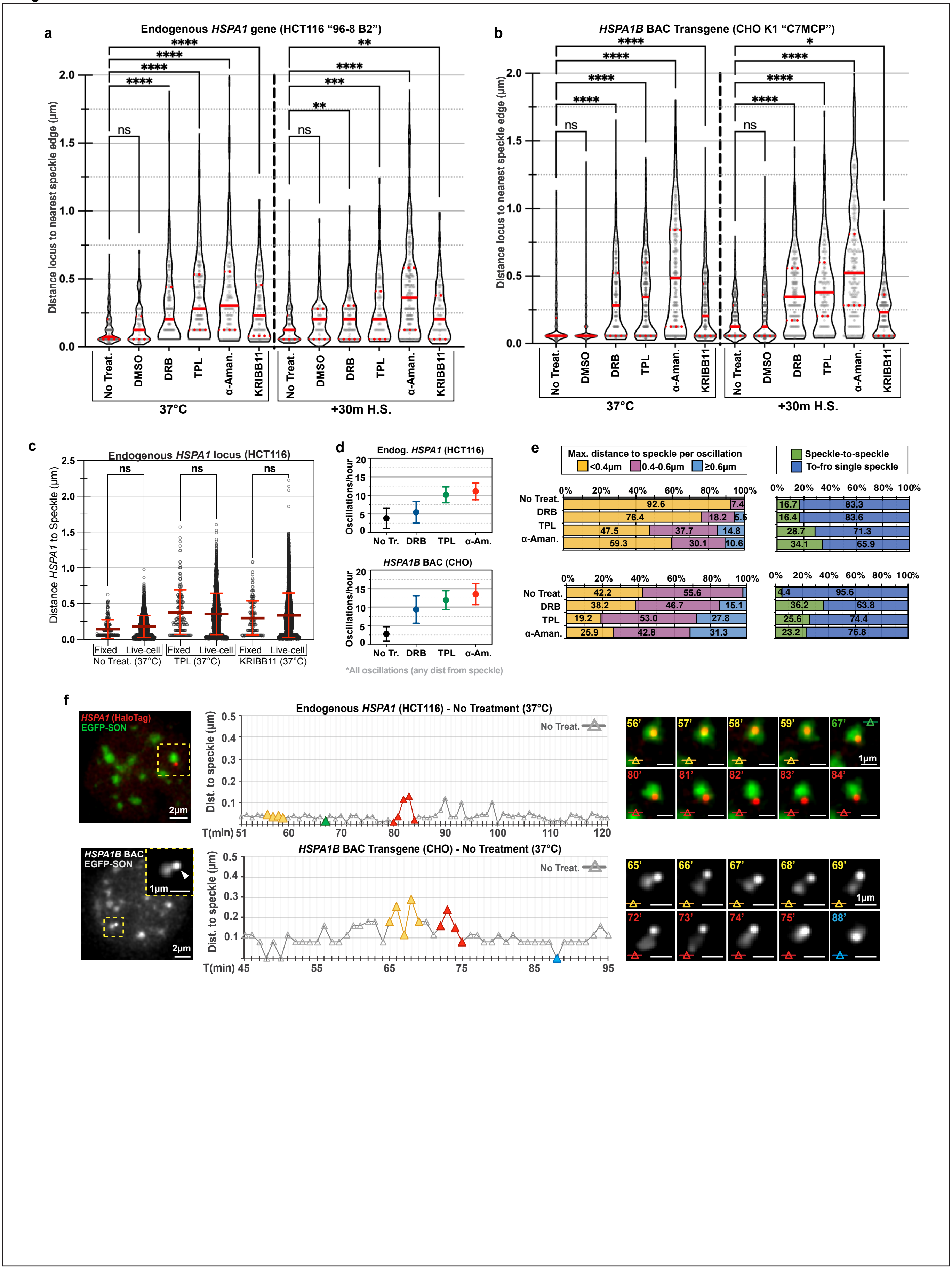

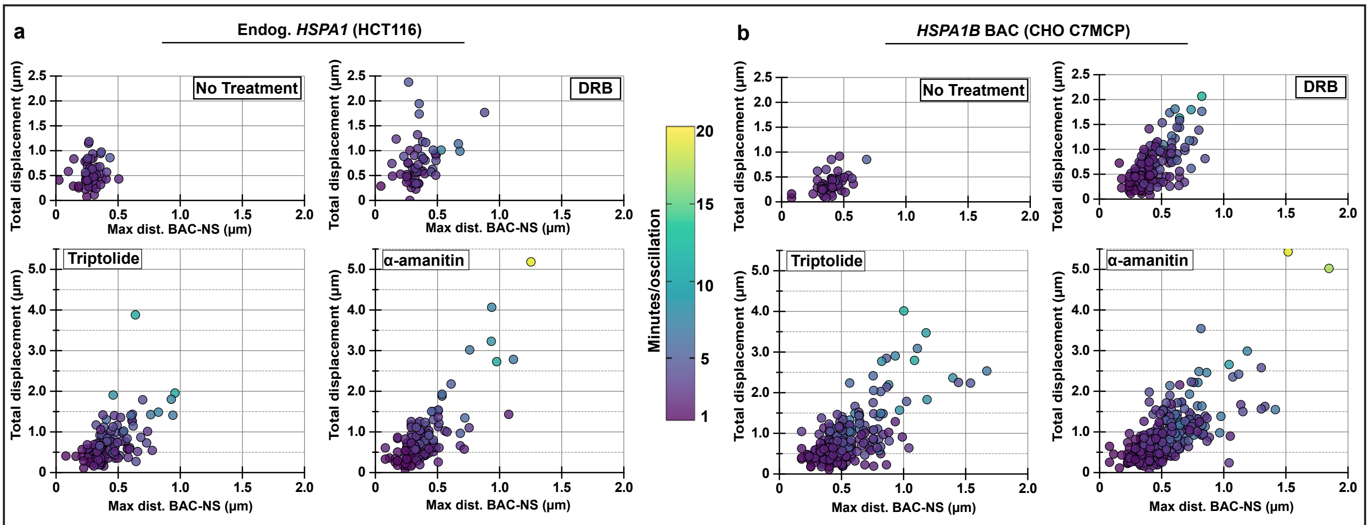

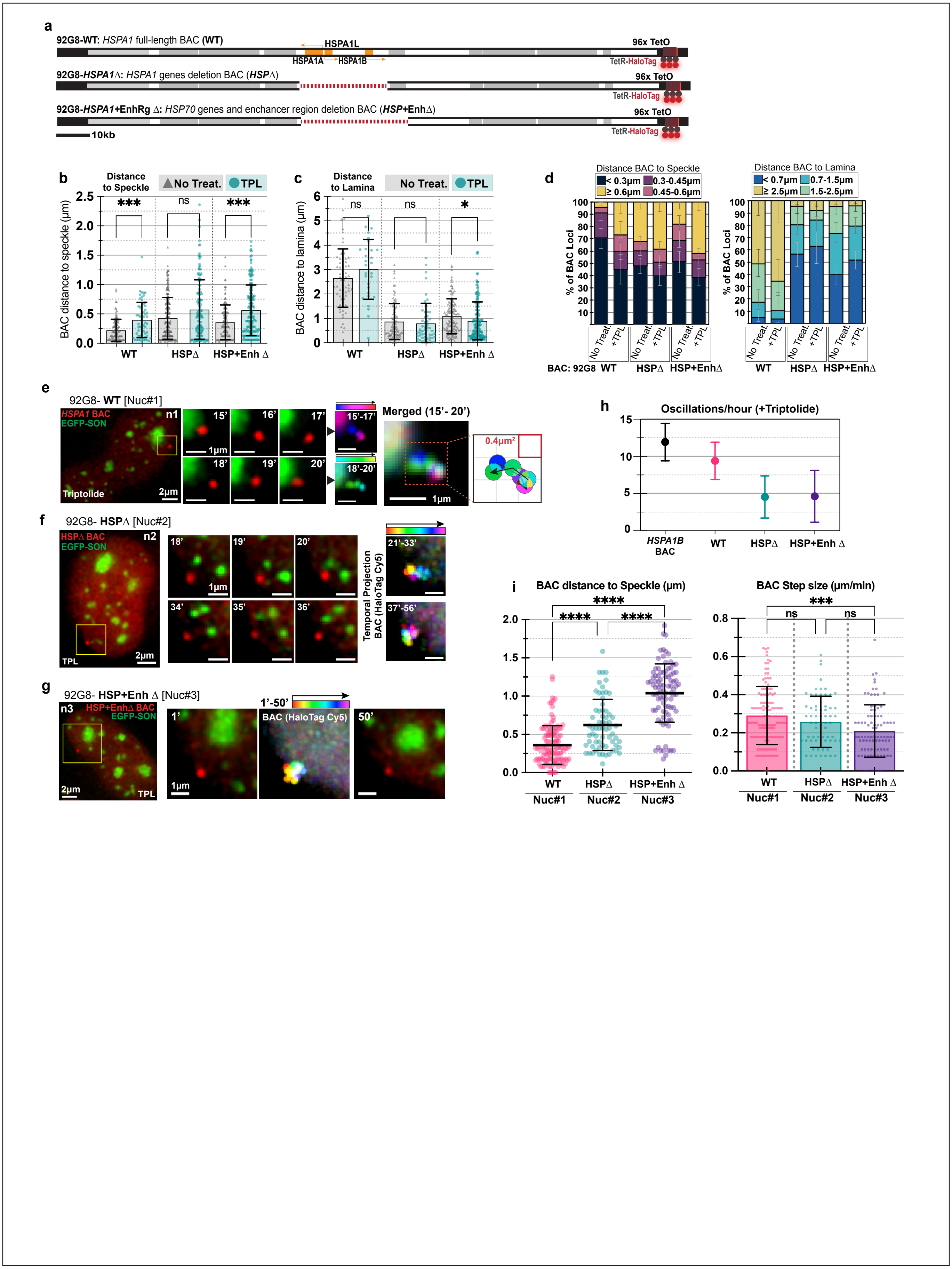

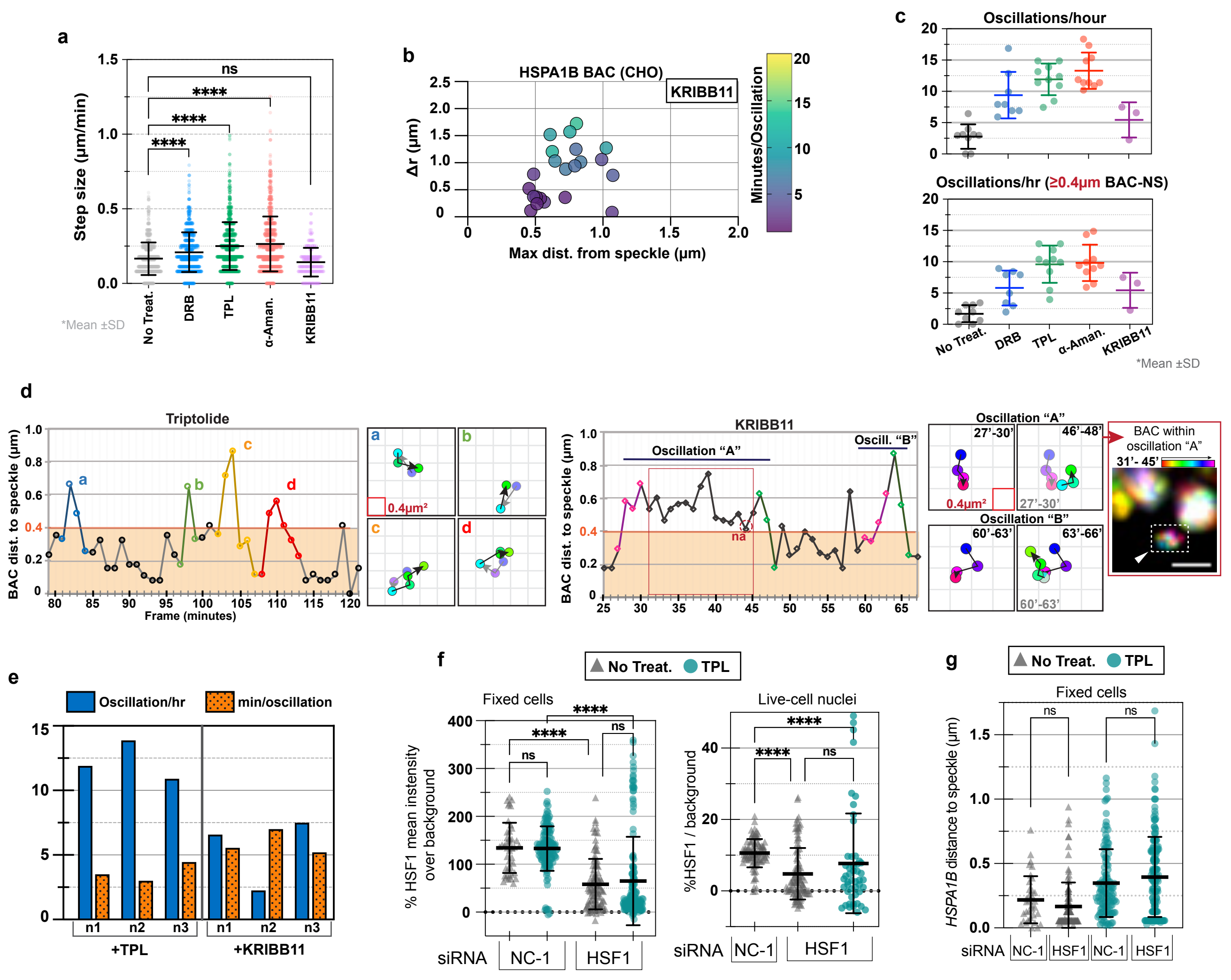

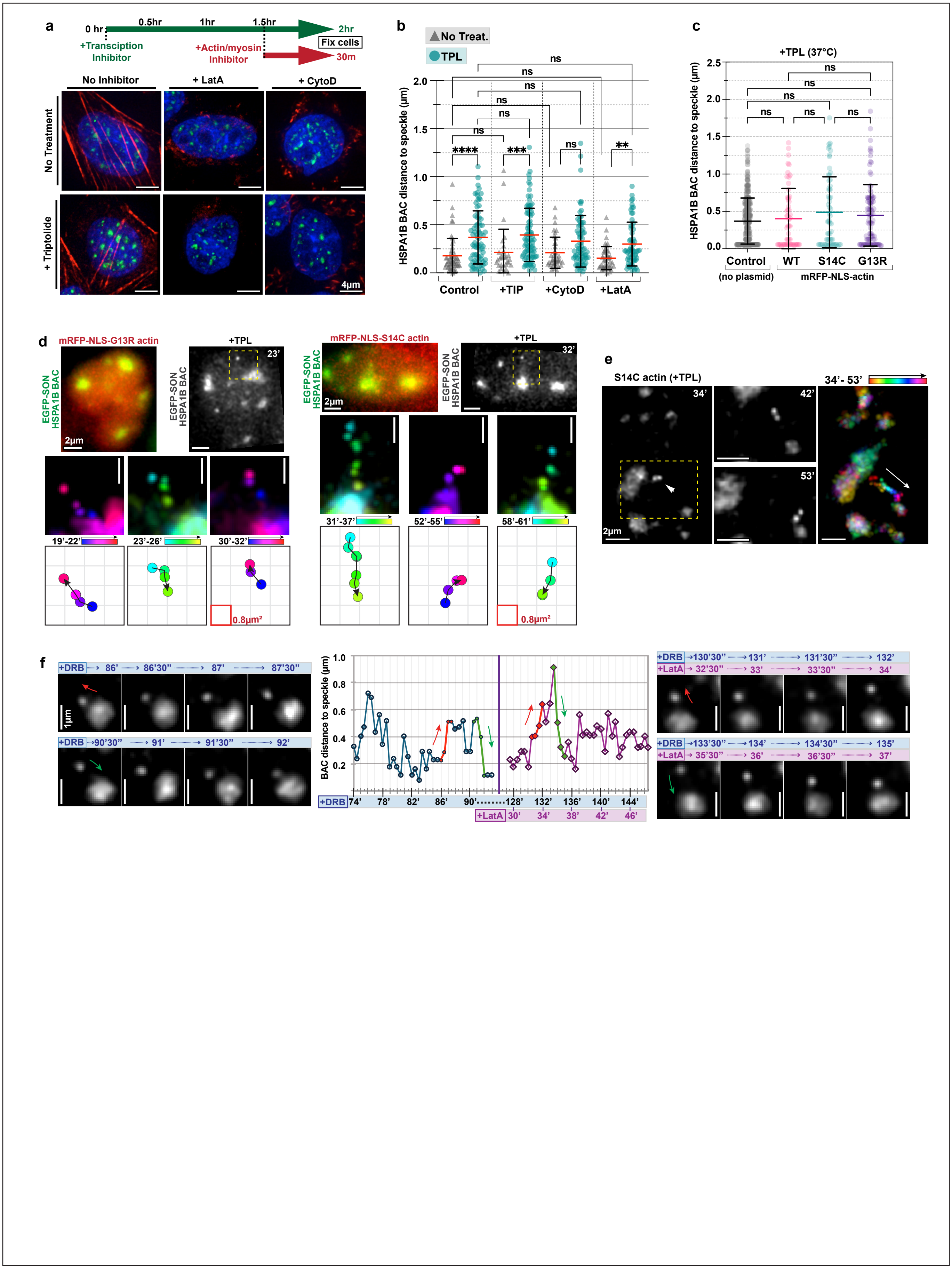

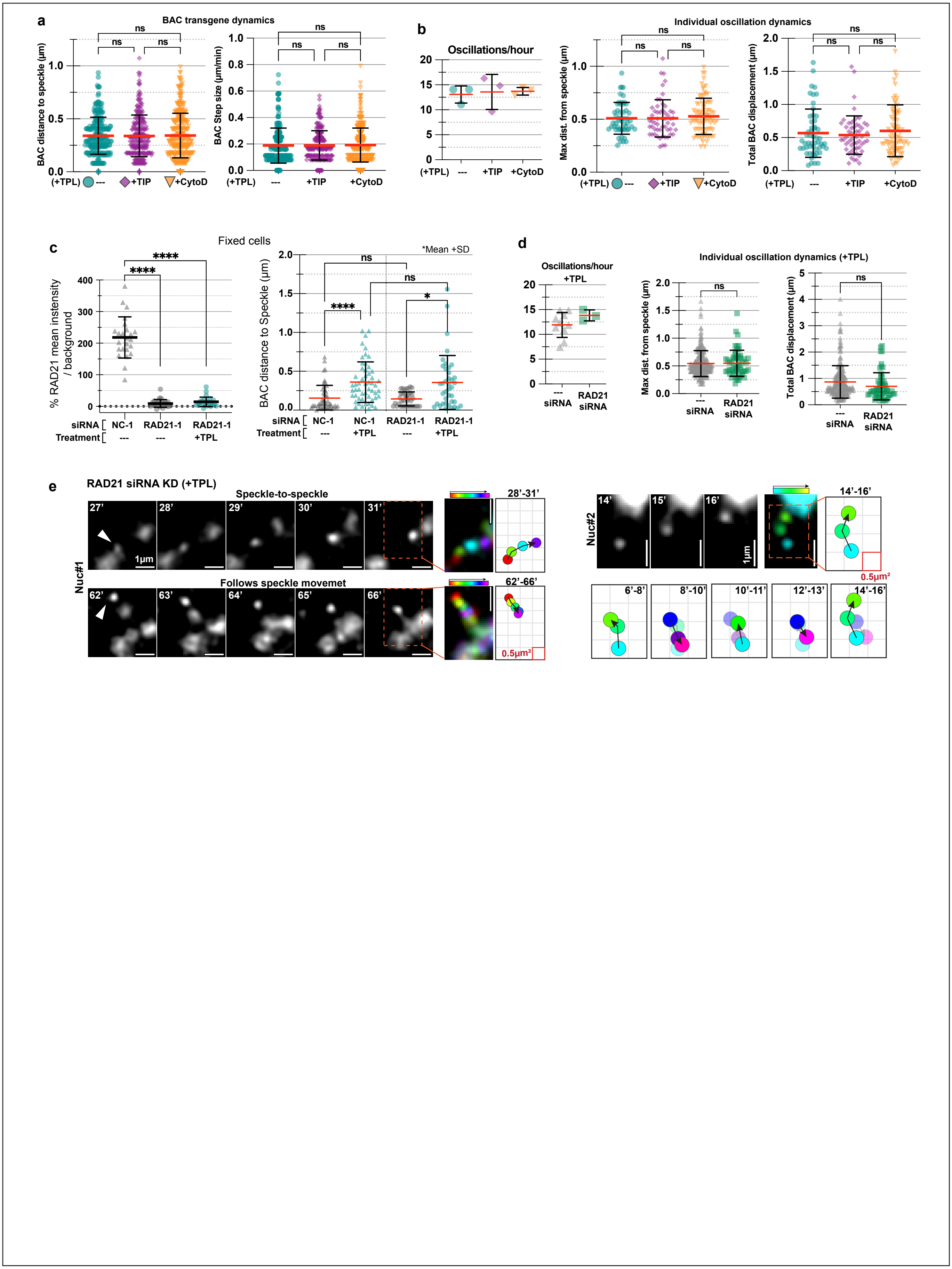

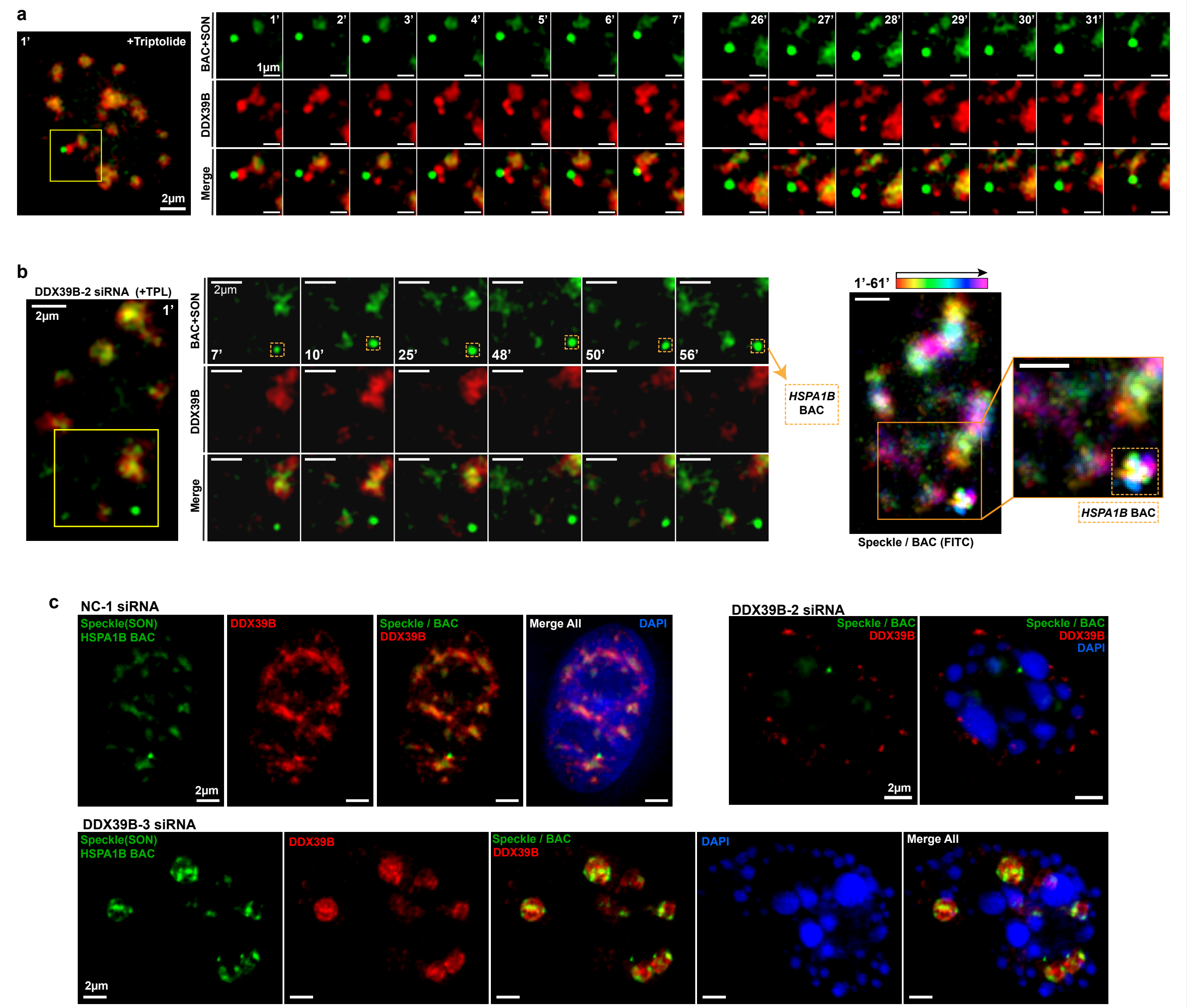
