## Supplementary Table 1 for "HSF1-dependent, long-range directional *HSPA1* gene motion to nuclear speckles is coupled to DDX39B condensate dynamics"

| **Generated Cell Line / Construct / Experiment** | **Target Locus** | **Component Type** | **Component Name** | **Sequence (5'-3')** |
| --- | --- | --- | --- | --- |
| **HCT116 96-8 B2** | *SON* | sgRNA | hSON-gRNA-1 | GAGAGAACGGAGCGGACGCCA |
|  |  | Homology Arm (5') | Left HA (Upstream) | GTGCTCACTGATTGG...AGCGGACGCC* |
|  |  | Homology Arm (3') | Right HA (Downstream) | GCGACCAACATCGAG...GCAGTCGTTC* |
| **CHO C7MCP HaloTag-HSF1 2D5** | *HSF1* | sgRNA | HSF1-gRNA-2 | TCCACCCGCTTCGTCCGAG |
|  |  | 5'-HA PCR primer (Fwd) | CHO_HSF1halo_LH | GACGACACTAGCTCA...TGGCACAGGA* |
|  |  | 3'-HA PCR primer (Rev) | CHO_HSF1halo_RH | AGCTTGGTCAGGAAG...TGAGCCAGAG* |
| **CHO C7MCP HaloTag-DDX39B 3D4** | *DDX39B* | sgRNA | DDX39B-gRNA-3 | ATTCTCTGCCATAATTGGGC |
|  |  | 5'-HA PCR primer (Fwd) | CHO_DDXhalo_LH | CCCCTTGCTTTCTCA...TGGCACAGGA* |
|  |  | 3'-HA PCR primer (Rev) | CHO_DDXhalo_RH | GCTGCTGTCTCCACC...TGAGCCAGAG |
| **CHO C7MCP HaloTag-RAD21** | *RAD21* | sgRNA | RAD21-gRNA-1 | CCAGAGGCCCTCTTTTACTG |
|  |  | 5'-HA PCR primer (Fwd) | CHO_Rad21_LH | TCCTCACATGAAACT...TTGGCACAGG* |
|  |  | 3'-HA PCR primer (Rev) | CHO_Rad21_RH | ATGGGCCGCCAGCCA...CTGAGCCAGA* |
| **RP11-92G8-RFTD** | RP11-92G8 | Homology Arm | 5' HR (TetO insertion) | GTCTCATTTTCGCCA...TGCAGTTTGC* |
|  |  | Homology Arm | 3' HR (TetO insertion) | GATGCCGGAGTCTGA...AGTGCTCGCC* |
| **RP11-92G8-RFTD-D3** | *HSPA1* | Recombineering Primer | *HSPA1* deletion Fwd (*galK*) | CACCATAATCCCATC...TCCTGTTGAC* |
|  |  | Recombineering Primer | *HSPA1* deletion Rev (*galK*) | TATTTTCTCCATGTA...GCTAGTGCGT* |
|  | *galK* | Recombineering Primer | *galK* removal Fwd | GTGGACTCCGCTCAC...TTCAGCATCT* |
|  |  | Recombineering Primer | *galK* removal Rev | TTCTCCATGTAATGG...GAATCTCCTG* |
| **RP11-92G8-RFTD-D3B (HSseD)** | *galK* + SE | Recombineering Primer | Enh + *galK* removal Fwd | TGTACCATTGTTCTG...CTCTTGACAA* |
|  |  | Recombineering Primer | Enh + *galK* removal Rev | TTCATGGACTGAGAC...TTGTTATAGA* |
| **siRNA Knockdown (CHO)** | *HSF1* | siRNA | HSF1-3 | UGAAGAGUGAGGACAUAAAAAUACG |
|  |  | siRNA | HSF1-4 | AACAAGCUCAUUCAGUUCCUGAUCT |
|  | *DDX39B* | siRNA | DDX39B-2 | GUCAUGAUGUUCAGUGCUACCUUGA |
|  |  | siRNA | DDX39B-3 | AUGAGCUCUUGGACUAUGAAGACGA |
|  | *RAD21* | siRNA | RAD21-1 | CCAUCACUCUACCUGAAGAAUUCCA |
|  |  | siRNA | RAD21-2 | GCUCAGUGAUUAUUCGGAUAUUGTG |
| *Full sequences are provided in the accompanying Extended Data Table 1 source file. | | | |  |

**Extended Data Table 1 | Oligonucleotides and reagents used for genome engineering.**

List of CRISPR-Cas9 sgRNAs and corresponding homology arm sequences or PCR amplification primers for donor DNA used to generate HaloTag endogenously tagged N-terminus cell lines (SON, HSF1, DDX39B, RAD21). Homology arm sequences and recombineering primers used for the modification of bacterial artificial chromosomes (BACs), including TetO array insertion and two-step gene deletions (HSPA1, galK), are listed. siRNA sequences used for transient knockdown experiments are provided in the final section. All sequences are provided in the 5' to 3' orientation. Full homology arm and primer sequences are truncated in the table; complete sequences are available in the accompanying Extended Data Table 1 source file.
